# NPOmix: a machine learning classifier to connect mass spectrometry fragmentation data to biosynthetic gene clusters

**DOI:** 10.1101/2021.10.05.463235

**Authors:** Tiago F. Leão, Mingxun Wang, Ricardo da Silva, Alexey Gurevich, Anelize Bauermeister, Paulo Wender P. Gomes, Asker Brejnrod, Evgenia Glukhov, Allegra T. Aron, Joris J. R. Louwen, Hyun Woo Kim, Raphael Reher, Marli F. Fiore, Justin J.J. van der Hooft, Lena Gerwick, William H. Gerwick, Nuno Bandeira, Pieter C. Dorrestein

**Author notes:** to whom correspondence should be addressed regarding the creation and management of the training data (Justin J.J. van der Hooft). **Author Contribution:** T.F.L. conceptualized the software; T.F.L., R.d.S., and A.B (Asker Brejnrod) programmed the software; M.W. and A.G. assembled the metagenomic reads and annotated all biosynthetic gene clusters; A.B. (Anelize Bauermeister) and R.R. worked on the predicted fragmentation for brasilicardin A; P.W.P.G. matched the predicted structure to the fragmentation of promising candidates and created structural insights; T.F.L, J.J.J.v.d.H., and M.W. curated the dataset and verified the known BGCs-metabolites links; T.F.L, J.J.R.L., J.J.J.v.d.H., and A.G. compared the performance of our tool with other multi-omics tools; E.G. cultured cyanobacterial samples and collected the cyanobacterial LC-MS/MS data published at PoDP database; J.J.J.v.d.H., H.W.K., R.R., A.T.A, P.W.P.G., and A.B. provided feedback on the approach and how to present the results; T.F.L and J.J.J.v.d.H. wrote the manuscript; J.J.J.v.d.H., L.G., W.H.G, N.B., M.F.F., and P.C.D. funded and designed the research; L.G., W.H.G, N.B., M.F.F., J.J.J.v.d.H, and P.C.D. edited the manuscript; all authors read, reviewed, and agreed to the published version of the manuscript.

## Abstract

Microbial specialized metabolites are an important source of and inspiration for many pharmaceutical, biotechnological products and play key roles in ecological processes. However, most bioactivity-guided isolation and identification methods widely employed in metabolite discovery programs do not explore the full biosynthetic potential of an organism. Untargeted metabolomics using liquid chromatography coupled with tandem mass spectrometry is an efficient technique to access metabolites from fractions and even environmental crude extracts. Nevertheless, metabolomics is limited in predicting structures or bioactivities for cryptic metabolites. Linking the biosynthetic potential inferred from (meta)genomics to the specialized metabolome would accelerate drug discovery programs. Here, we present a *k*-nearest neighbor classifier to systematically connect mass spectrometry fragmentation spectra to their corresponding biosynthetic gene clusters (independent of their chemical compound class). Our pipeline offers an efficient method to link biosynthetic genes to known, analogous, or cryptic metabolites that they encode for, as detected via mass spectrometry from bacterial cultures or environmental microbiomes. Using paired data sets that include validated genes-mass spectral links from the Paired Omics Data Platform, we demonstrate this approach by automatically linking 18 previously known mass spectra to their corresponding previously experimentally validated biosynthetic genes (i.e., via NMR or genetic engineering). Finally, we demonstrated that this new approach is a substantial step towards making *in silico* (and even *de novo*) structure predictions for peptidic metabolites and a glycosylated terpene. Altogether, we conclude that NPOmix minimizes the need for culturing and facilitates specialized metabolite isolation and structure elucidation based on integrative omics mining.

**Significance:** The pace of natural product discovery has remained relatively constant over the last two decades. At the same time, there is an urgent need to find new therapeutics to fight antibiotic-resistant bacteria, cancer, tropical parasites, pathogenic viruses, and other severe diseases. Here, we introduce a new machine learning algorithm that can efficiently connect metabolites to their biosynthetic genes. Our Natural Products Mixed Omics (NPOmix) tool provides access to genomic information for bioactivity, class, (partial) structure, and stereochemistry predictions to prioritize relevant metabolite products and facilitate their structural elucidation. Our approach can be applied to biosynthetic genes from bacteria (used in this study), fungi, algae, and plants where (meta)genomes are paired with corresponding mass fragmentation data.

## Introduction

Microbial specialized metabolites are often made by biosynthetic genes that are physically grouped into clusters known as biosynthetic gene clusters (BGCs). Currently, the most common drug discovery methods utilize metabolomics but do not make use of genomics and, consequently, they cannot explore the full biosynthetic potential found in nature. In fact, it has very recently been estimated that metabolomics could only discover 3% of the bacterial biosynthetic potential so far (1). This percentage indicates the number of classes in the NPAtlas structure database (2) that corresponds to the predicted potential of the bacterial kingdom (over a million gene clusters known for encoding specialized metabolites), estimated by Gavriilidou and collaborators, 2022 (1). Additionally, the great majority of these metabolomics approaches rely on culturing and/or chromatographic isolation for bioactivity assays and structure elucidation. Recent advances in computational metabolomics and metabolome mining do also facilitate specialized metabolite discovery and characterization (3). However, the full elucidation of a structure of a metabolite is usually done via a set of 1D/2D nuclear magnetic resonance (NMR) experiments or X-ray crystallography, two techniques that are expensive and time-consuming. Genome mining would allow making accurate *in silico* predictions about the structure, bioactivity, and novelty of a given unknown metabolite from microbial cultures or even environmental microbiomes.

One of the challenges in the genome mining field is to connect microbial metabolites to their BGCs with confidence. Even the genome of *Streptomyces coelicolor* A3(2), one of the first sequenced microbial genomes, still contains a large number of cryptic BGCs – BGCs without known associated metabolites (4). In 2011, the bioinformatics tool antiSMASH (5) improved the identification and annotation of BGCs based on automated genome mining. AntiSMASH annotates BGCs by searching them with profile hidden Markov models of domains from gene/protein sequences known to biosynthesize metabolites and these models are specific to a certain class of BGC. We used antiSMASH to annotate all BGCs in this study. Moreover, since 2018, BiG-SCAPE (6) can reliably calculate the similarity between pairs of BGCs, grouping them into gene cluster families (GCFs). Recently, some approaches and tools have been created to connect specialized metabolites (known and cryptic MS/MS spectra) to their biosynthetic gene clusters, such as Pattern-based Genome Mining (7, 8), MetaMiner (9), DeepRiPP (10), NRPquest (11), NRPminer (12), GNP (13) and NPLinker (14), recently reviewed by Van der Hooft *et al*., 2020 (15). Nerpa (16) and GARLIC (17) can connect structures to BGCs; structures are normally represented in the SMILES (Simplified Molecular-Input Line-Entry System) format, a type of computer-readable annotation language for chemical structures. However, most of these tools are neither high throughput, nor efficient, or can only be used for a particular class of BGC (e.g., peptides or BGCs homologous to known BGCs). It has been challenging to create a systematic tool that can work at the repository scale to connect genotypes (BGCs) with their phenotypes (for example fragmentation spectra, MS/MS spectra, from untargeted liquid chromatography coupled with mass spectrometry profiles, LC-MS/MS). As a result, a large disparity exists between the number of known metabolites from an organism versus the number of BGCs with known metabolite products. For example, the recently designated cyanobacterial genus *Moorena* has already yielded over 200 metabolites, yet only a dozen of validated BGCs are currently deposited for this genus in the expert-annotated Minimum Information about a Biosynthetic Gene cluster (MIBiG) database (18). Connecting metabolites to their biosynthetic genes would also facilitate research concerning the ecological role and functions of the specialized metabolome by studying the regulation of the expression of their biosynthetic gene clusters.

As an analogy, structure prediction for proteins is extremely challenging and a brute force approach would take longer than the age of the universe to correctly predict a structure of a single protein, an issue that was named “the protein structure problem”. A similar problem exists for metabolite discovery and these structures are key for mining novel and bioactive specialized drug-like candidates (via docking or specific genes that already can be used to indicate bioactivity). The deep learning algorithm named AlphaFold 2 (19) was able to use only amino acid sequences to make protein structure predictions with over 90% precision and this precision was considered (by the comparative modeling community) as enough to solve “the protein structure problem”. A part of the reason this problem could be solved was because billions of dollars were spent in the past few decades to solve protein structures, whereas, in contrast, training sets for genomes, metabolomes, and metabolite structure links are just becoming available. So, one of the main questions is how we can begin solving the “metabolite structure problem” and what algorithms are needed to do so? It is certainly a very challenging task, but we believe that the first step toward it is to use the power of genomics for *in silico* predictions by developing a tool that can efficiently connect known or cryptic metabolites (detected via LC-MS/MS) to their correct BGC.

*In silico* predictions using raw DNA sequence and unlabeled mass fragmentation data can be used to predict: bioactivity, either by gene content or docking experiments; full planar structures, by dereplication, homology, or *de novo*; partial stereochemistry, and; novelty, based on the abundance from all bacteria sampled from nature so far. For example, some random forest classifiers by the Clardy lab (20) at Harvard medical center use only the BGC sequences to predict with about 80% precision if a BGC will produce an anticancer, antifungal, or antibacterial metabolite. Coelichelin was isolated using an *in silico* structure prediction that indicated the peptide to be a siderophore, which was confirmed by culturing the producer in an iron-deficient media and isolating the induced metabolite (overexpressed when compared to the control). BiG-SLICE was used to predict the connections between more than 1 million BGCs, providing clues to the most unique BGCs in thousands of samples.

To start addressing the above-mentioned “metabolite structure problem” and the gap between genomics and metabolomics in drug discovery, we deployed a *K*-Nearest Neighbor (KNN) algorithm that uses similarity BGC fingerprints and analogously similarity MS/MS fingerprints to classify gene cluster family (GCF, a group of similar BGCs) candidates for each MS/MS spectrum. We showed that the addition of biosynthetic class and substructure features improves the performance of our Natural Products Mixed Omics tool (NPOmix, available at https://www.tfleao.com/npomix1). While our approach uses unique fingerprints and machine learning algorithms for connecting metabolites to BGCs, it can be considered a type of Pattern-based Genome Mining which was previously reported by Doroghazi *et al.* in 2014 and Duncan *et al.* in 2015 (7, 8). Pattern-based Genome Mining is based on the idea that the distribution of a given natural product should be comparable to the distribution of the BGCs responsible for their production. We would like to acknowledge that this approach is limited to organisms that have BGCs (not all organisms have BGCs as well-defined as bacteria and fungi, for example, higher plants and animals typically have less well-structured clusters of biosynthesis genes) and to BGCs that are somewhat similar to at least one of the reference BGCs in the Paired Omics Data Platform (PoDP)(21) datasets used by NPOmix. Benchmarked to eight other multi-omics tools, NPOmix is the only currently available tool that is: i) scalable – it can run on a personal laptop with 8GB of RAM and four cores for 550 metabolites in less than 4 hours, using over four thousand genomic/metabolomic files; ii) systematic – it works for many different NP classes (e.g., non-ribosomal peptides, polyketides, terpenes, and so on); and iii) highly efficient – its maximum performance was markedly higher than the other tested tools with a 92.3% precision during benchmarking (using a single nearest neighbor candidate and 22 validated BGCs-metabolite links). Of note, in this study, six of the 11 BGC-metabolite links used for validation (not counting analogs) were previously fully characterized via knockouts, heterologous expression, and/or isolation, and unambiguous NMR structure elucidation as recorded at the PoDP. The number of genomes, metabolomes, BGCs, GCFs, GNPS metabolites, metabolites with MIBiG BGC, and BGC-metabolite links (with and without filtering) are listed in Table S1. The major limitation of the evaluation of our method was the lack of available test data for structures that are linked to their MS/MS spectra and biosynthetic gene clusters - a bottleneck that the application of NPOmix can help to solve and we discuss more details in the Supplementary Information, Background section. We believe that our NPOmix tool will assist with the discovery of novel metabolites as well as known metabolites with new biosynthesis (more details in the Supplementary Information Background).

## Results

### Our proposed solution: the Natural Products Mixed Omics (NPOmix) approach

To enable *in silico* predictions of molecular novelty, bioactivity, and chemical structure, metabolites need to be connected to their corresponding BGCs, a major challenge in the metabolite discovery field that has been tackled by eight different approaches that are: MetaMiner (9), DeepRiPP (10), NRPquest (11), NRPminer (12), GNP (13) and NPLinker (14)(that includes Metcalf’s co-occurrence-based Pattern-based Genome Mining and a standardized form thereof)(7, 8), Nerpa (16) and GARLIC (17). However, i) none of these approaches are widely used for drug discovery, and ii) genomics is often not included in drug discovery pipelines, despite the numerous advantages of using paired genomics-metabolomics for *in silico* predictions (as exemplified in the introduction). Therefore, we decided to leverage the power of multidisciplinary expert knowledge (from a team including 2 molecular biology, 11 metabolomics, 1 genomics, and 5 multi-omics experienced researchers from around the globe) and machine learning to create a high-performance KNN classifier to solve this issue. Our KNN classifier named Natural Products Mixed Omics (NPOmix) can use just DNA sequences and unlabeled mass fragmentation profiles to accurately create links between metabolites and their corresponding biosynthetic gene cluster.

To use the NPOmix approach, it is required to have a paired dataset with genomic and MS/MS information. Figure 1 shows a conceptual example using only four samples and only using similarity as a feature. The genomic information can be either that of a genome or metagenome and the MS/MS spectra should be obtained via untargeted LC-MS/MS. Paired datasets have become available at the Paired Omics Data Platform (PoDP)(21), one of the first initiatives to gather paired genomic and MS/MS information. Our analysis contained 3,331 BGCs (used for training) that were present in 1,040 networked genomes/metagenomes with paired LC-MS/MS data that could be downloaded from the PoDP database.

**Fig. 1.**
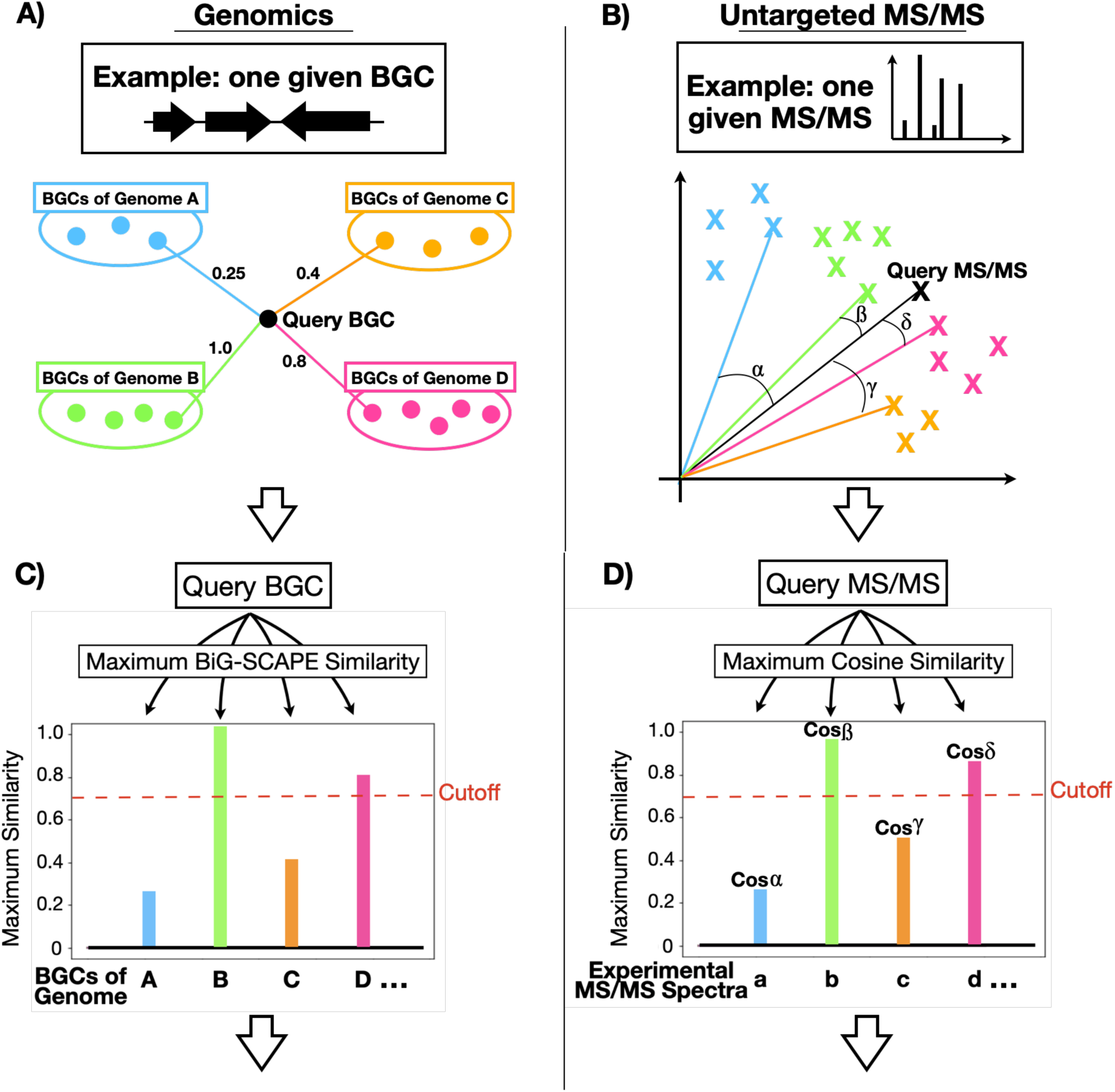
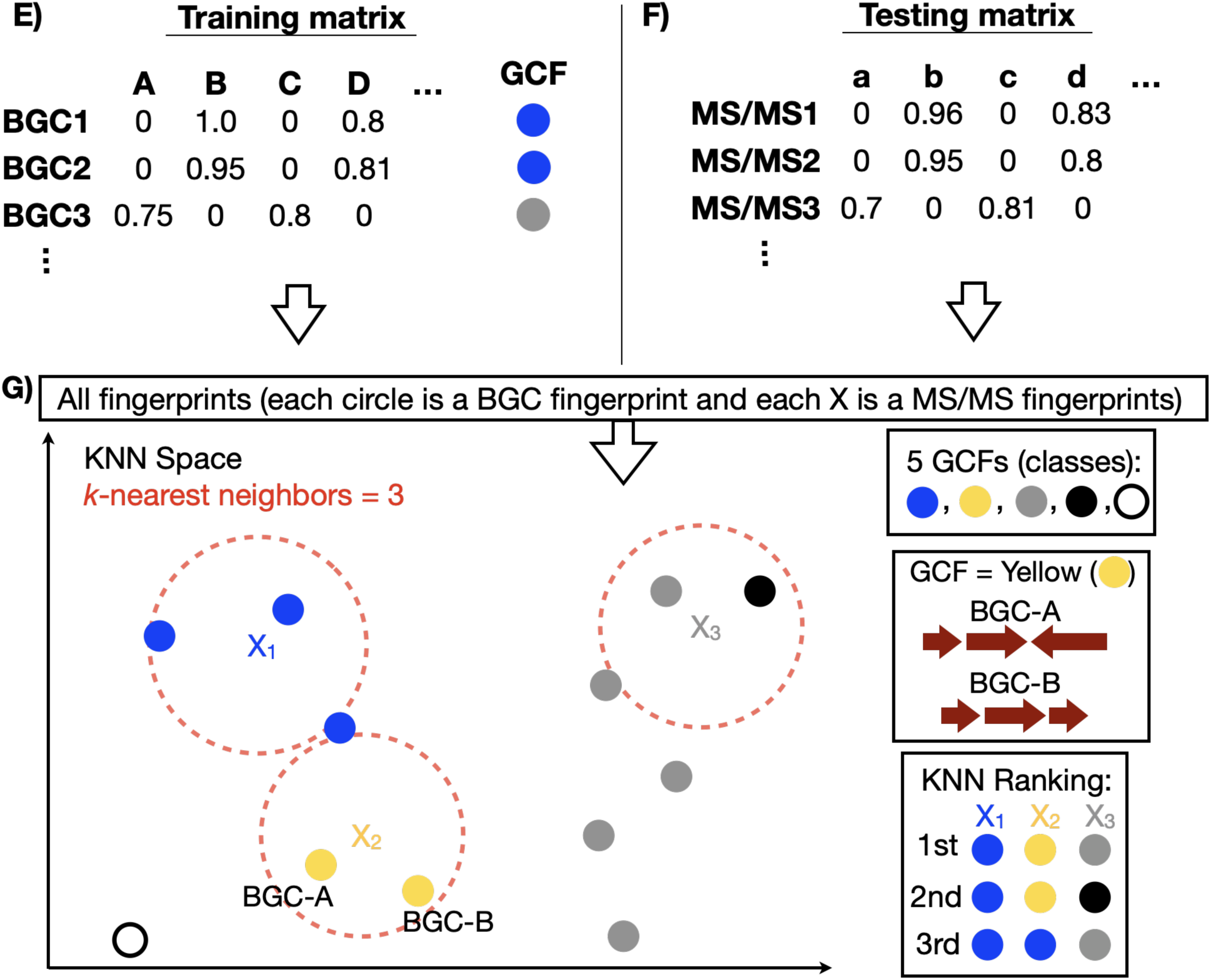
The genomics and metabolomics pipelines use the proposed KNN approach for a hypothetical dataset with 4 paired genomes-MS/MS samples. Representation of how to calculate the similarity scores between BGCs (displayed in A) and between MS/MS spectra (displayed in B). Schematic of how to create BGCs (C) and MS/MS (D) fingerprints using a paired genomics-metabolomics dataset of four samples, i.e., genomes, metagenomes, or MAGs (samples A-D) and similarity scores from BiG-SCAPE and GNPS. The dashed red line represents the selected cutoff of 0.7. The query BGC is highly similar to a BGC in sample B (indicating an identical BGC), while it is probably absent in samples A and C. The BGC fingerprints are grouped together in a training matrix and the MS/MS fingerprints compose the testing matrix (F). All fingerprints are plotted in the multi-dimensional KNN space (G, here represented in only 2D for simplification) where each circle represents a BGC fingerprint, and each X represents an MS/MS fingerprint. BGCs are labeled according to one of the five GCFs (five differently colored circles). KNN ranking of neighbors is based on the proximity between the query MS/MS fingerprint and the neighboring BGC fingerprints. In this example, a KNN = 3 (three closest neighbors) is depicted. BGC = biosynthetic gene cluster; MS/MS = mass fragmentation spectrum; KNN = *K*-nearest neighbor; BiG-SCAPE = software to calculate pairwise BGC-BGC similarity; GCF = gene cluster family (group of similar BGCs); Cosine score = modified cosine score from GNPS to calculate pairwise spectrum-spectrum similarity. For example, the blue circle shared between X1 and X2 could represent a lipopeptide that shares the peptidic portion with other blue circles and a fatty acid portion with the yellow circles or vice versa.

We first used grouped known and cryptic BGCs in GCFs based on domain similarity using BiG-SCAPE (6) to obtain labels that can be used for supervised learning (Fig. 1A). To create a BGC fingerprint (Fig. 1C), we identified the similarity between the query BGC and each of the BGCs in each genome in the training dataset (similarity scores are computed as “1 minus the raw distance”). The BGC fingerprint that emerges is a series of columns for each compared genome, the column value of which represents the similarity score between the query BGC and the BGC to which it is maximally similar in each genome (column). Similarity scores range from 0.0 to 1.0; identical BGCs have perfect similarity and are scored as 1.0 whereas a score of 0.8 would indicate that a homologous BGC is present in the genome. A score below the (user-defined) similarity cutoff of 0.7 indicates that the queried BGC is likely absent in the genome. We selected the cutoff as 0.7 because it is the same cutoff used for BiG-SCAPE (6) similarity (a value which was determined using a cutoff calibration with the MIBiG database)(22). An analogous process is used to create MS/MS fingerprints (Fig. 1B and 1D); a query MS/MS spectrum is compared to all of the MS/MS spectra in the query set. Furthermore, we add the presence/absence of biosynthetic classes and substructures to the BGC and MS/MS fingerprints (for more details, refer to the Methods section). We then use the BGC fingerprint in the training together with the generated family labels. In more detail, the BGC fingerprints form a training matrix (Fig. 1E, where each column is a genome and each value is the maximum similarity between the query BGC and the BGCs in this given genome) and, in this case, the matrix contains 1,040 columns due to the 1,040 sets of paired experimental samples plus the columns corresponding to the biosynthetic class and annotated substructures.

The trained KNN classifier takes as input the same kind of features (MS/MS fingerprint, Fig. 1D, and the testing matrix is exemplified in Fig. 1F, where the similarity is now represented by the modified cosine score) but these features are annotated from individual mass fragmentation profiles (known or cryptic metabolites). The aim of NPOmix is then to label these MS/MS fingerprints with the proper gene cluster family, thus accurately linking metabolites to their corresponding BGCs. The query MS/MS spectrum could be either a reference spectrum from the Global Natural Products Social Molecular Networking database (GNPS) (23, 24) or a cryptic MS/MS spectrum from a new sample that contains a sequenced genome and experimental MS/MS spectra. In more detail, the KNN algorithm links BGCs to metabolites by plotting the BGC fingerprints in the KNN feature space (in Fig. 1G). The KNN feature space is exemplified by only two dimensions using dimension reduction techniques since a 1,040-dimensional space is not feasible to comprehensively visualize (one dimension per sample).

More details of how this multidimensional plotting occurs are illustrated in Fig. S1, where the samples correspond to the 3D axes and the similarity scores are the coordinates. Each query MS/MS fingerprint (a row in the testing metabolomic matrix and columns are the experimental MS/MS spectra per sample) is plotted into the same KNN feature space (Fig. 1G) so the algorithm can obtain the GCF labels for the nearest neighbors to the query MS/MS fingerprint (e.g., for three most similar BGC neighbors, *k* = 3). We note that GCF labels can be present more than once in the returned list if two or more BGC nearest neighbors belong to the same GCF. Given that many related BGCs are part of the same GCF, this repetition of the GCF classification is a common behavior of our KNN approach.

In principle, our approach is suitable for bacterial (exemplified here), fungal, algal, and plant genomes and MS/MS spectra obtained from the same organism (if these organisms contain BGCs). Metagenomes and metagenome-assembled genomes (MAGs) can also be used instead of genomes; however, complete genomes are preferred in the training set primarily due to the expected data quality. This KNN approach also supports LC-MS/MS from fractions or different culture conditions; multiple LC-MS/MS files for the same genome were merged into a single set of experimental MS/MS spectra.

### Validation and multi-omics dereplication: linking known metabolites to known BGCs, validated links from the PoDP database and BGCs from the MIBiG database

To validate our NPOmix approach, we used 36 out of 71 datasets from the Paired Omics Data Platform (PoDP, from February 2021, datasets listed at Dataset S1, sheet one). There are currently 75 datasets in the PoDP database. We selected genomic samples that contained a valid Genome ID or BioSample ID to aid their downloading from the National Center for Biotechnology Information database and totaling 732 genomes/MAGs obtained from these 36 meta-datasets. We also selected and assembled 1,034 metagenomes from two major metagenomic datasets: 1) MSV000082969 and PoDP ID cd327ceb-f92b-4cd3-a545-39d29c602b6b.1 – 556 cheetah fecal samples and environmental samples (23); 2) MSV000080179 and PoDP ID 50f9540c-9c9c-44e6-956c-87eabc960d7b.3 – The American Gut Project (24) that contains fecal samples from 481 human subjects. These (meta)genomes were downloaded with the code shared at the GitHub repository https://github.com/tiagolbiotech/NPOmix, notebook 1. The LC-MS/MS files can be downloaded using “FTP” from links recorded in Dataset 1, sheet two. We were able to network 1,040 (meta)genomes that contained 3,331 BGCs (including 260 BGCs from the MIBiG database) distributed into 997 GCFs. In the untargeted metabolomics data, we matched 3,248 LC-MS/MS files to 22 GNPS (13, 25) reference library MS/MS spectra and one spectrum not available at GNPS (brasilicardin A, obtained from the microbial metabolome) to create the MS/MS fingerprints for testing the KNN classification (one fingerprint per spectrum). We envision to create a more balanced, diverse, and less sparse training dataset in ongoing efforts. To maximize precision rates in the future, we plan to purchase cultures from collections that have well-assembled genomes so we can obtain the paired LC-MS/MS. However, the current dataset produced highly supportive results by testing validated links from the PoDP, links generated by the Gerwick lab dataset, and semi-manually connected links used in the NPLinker publication, all of which will be reused for posterior benchmarking of BGC-MS/MS linking algorithms. We attempted to test all 242 BGC-metabolite links used as training data for the Rosetta scoring method in NPLinker (totaling 2,069 unique MS/MS spectra, Dataset S1, sheet two) plus 109 manually added MS/MS links (connected to BGCs, annotated by experts at the PoDP, Dataset S1, sheet three). However, most of these validated MS/MS spectra were not present in the 1,040 paired (meta)genomes-MS/MS samples from the PoDP or their BGC scores did not co-occur with their MS/MS scores because they were not present in the nearly exact same samples. To further illustrate this, 178 MIBiG BGCs networked with PoDP BGCs and 50 GNPS-annotated metabolites were reported at PoDP. The intersection of these two resulted in 22 BGC-metabolite links including analogs. You can see these and other metrics in Table S1.

Hence, our validation dataset was limited to 11 validated links (22 with analogs) from the MIBiG and PoDP databases and found in the paired (meta)genomes-MS/MS samples (barbamides, antimycin, pyocyanines, 2,4-diacetylphloroglucinol, brasilicardin A, orfamides, albicidins, bafilomycin B1, nevaltophin D, jamaicamides and cryptomaldamide, totaling 22 references MS/MS spectra that were present in the GNPS database). We only included analogs for which the MIBiG BGC was present in our paired samples downloaded from the PoDP; hence, we included orfamide A-C but not orfamide E, F and so on (because they are not reported as products from the MIBiG orfamide BGCs). We stress that a larger training dataset with more complete genomes is likely to increase the size of the validation set by adding more valid BGCs into the analysis. We were also able to combine NPOmix with *in silico* metabolomics tools like Dereplicator+ (20) to make new links between MS/MS spectra, BGCs, and molecular structures, as we will exemplify by brasilicardin A. This was accomplished by annotating cryptic MS/MS spectra (without a GNPS library hit and therefore not present in the GNPS database) to known BGCs (found in the MIBiG database). Such new links could be confirmed experimentally to improve the size of the validation set, as well as to expand MS/MS databases by adding these cryptic spectra to them.

A two-dimensional comparison of both types of fingerprints (BGC and MS/MS) can be a proxy for distinguishing some true positives from false positives. As observed in Fig. S2, we can visualize a mismatch between the BGC fingerprints (one GCF) and the MS/MS fingerprint in the “reduced” KNN-space (represented schematically in only two dimensions), indicative of a possible false positive link. This GCF is dereplicated as the known metabolite, pyocyanin, and it was incorrectly associated with the metabolite 2,4-diacetylphloroglucinol, confirming the false positive (at *k* = 3). In contrast, Fig. 2 illustrates that 5 metabolites, albicidin (structure in Fig. 2A and MS/MS spectra in Fig. 2B) and four albicidin analogs, could be correctly assigned to their corresponding GCF that contains 2 BGCs (one from the MIBiG database and another from *Xanthomonas albilineans* GPE PC73, GenBank ID GCA_000087965.1). In this case, the BGC fingerprints match the MS/MS fingerprints (Fig. 2C and 2D). This observation of “co-occurrence” between the strains in the BGC fingerprint and MS/MS fingerprints can be quantified by the Jaccard index (a score ranging from zero to one and calculated by the intersection of the presence/absence between the strains in fingerprints over the total number of strains).

**Fig. 2.**
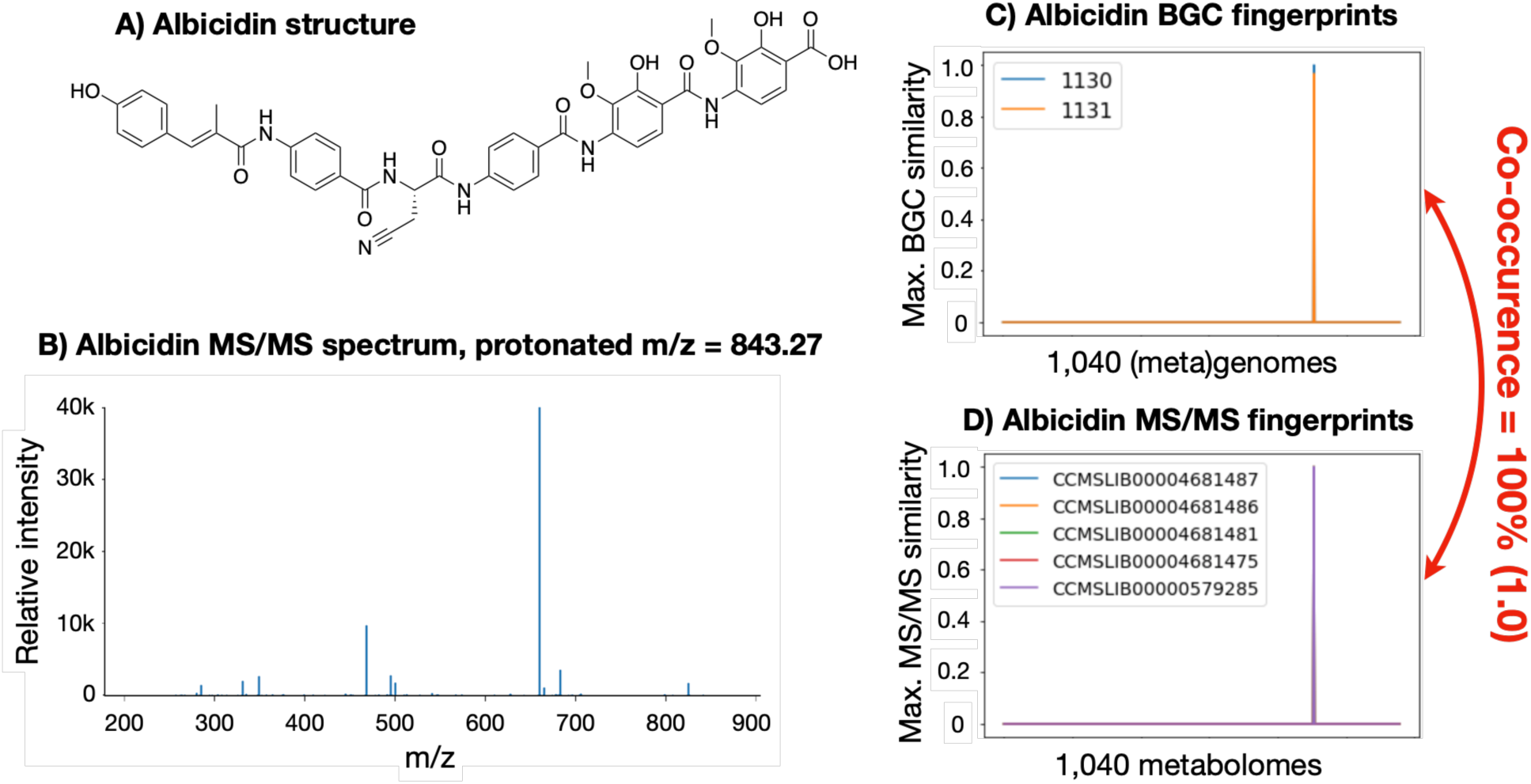
Multi-omics enabled the dereplication of albicidin by automatically predicting a true BGC-metabolite link. Structure of the dereplicated metabolite (A) and its corresponding representative MS/MS spectrum (B, spectrum example from GNPS ID CCMSLIB00000579285 and *m/z* of 843.2700), obtained via Metabolite Spectrum Resolver (26). The two BGC fingerprints (1130 and 1131) are represented in a 2D plot (C) and they match the 2D plot for the 5 MS/MS fingerprints obtained from GNPS for albicidin and its analogs (D). BGC = biosynthetic gene cluster; MS/MS = mass fragmentation spectrum; *m/z* = mass over charge calculated via mass spectrometry.

Using the PoDP dataset, a Jaccard index cutoff of 0.7 and similarity plus biosynthetic class as features for the machine learning model, we obtained a precision of 92.9% as 13 out of 14 references MS/MS spectra were correctly labeled when top-*n* = 3 (*k* nearest neighbors equal to 3, metabolites and their predicted GCFs listed in Table 1). Top-*n* represents how often the correct GCF label was found among the top *n* labels classified by the KNN approach. We have determined that the use of three neighbors is the optimal performance (also using the co-occurrence threshold), providing a good balance between precision and the number of links to validate (very high precision and randomness equal to 0, as detailed in Table S2). Randomness is observed by shuffling the testing columns, experimental MS/MS names, and counting how many correct links are present between the top-*n* GCF candidates. Lastly, we regard our NPOmix approach as multi-omics enabled dereplication (or *in silico* structure prediction by homology) because the 13 MS/MS spectra were automatically assigned to a known GCF that confirmed their metabolite labels, thereby minimizing the need for purchasing standards, performing isolation and NMR characterization, gene knockout or heterologous expression.

**Table 1.** 14 links between GNPS MS/MS spectra (with metabolite ID) and networked gene cluster family (true GCF that contained a known MIBiG BGC and passed the co-occurrence threshold). The table also includes their KNN predictions (*k* = 3); the predicted GCFs are ordered according to the value for *k*, from 1 (nearest) to 3 (furthest), and the first correct family is marked in bold red font. GCF labels can be repeated because multiple BGCs from the same GCF can be predicted as the nearest neighbors. Classification is considered correct if the true GCF is among the top-3 candidates. Annotations are according to each MIBiG BGC(s) found in the true GCFs. GNPS = Global Natural Products Social Molecular Networking database; KNN = *K*-nearest neighbor; MS/MS spectra = fragmentation spectra; BGC = biosynthetic gene cluster; GCFs = gene cluster family; MIBiG = Minimum Information about a Biosynthetic Gene cluster database; * = new analogous MS/MS spectra that were automatically connected to validated BGCs.

| Metabolite ID | True GCF | Predicted GCFs for $k = 3$ | Annotation |
| --- | --- | --- | --- |
| CCMSLIB00004679298 | GCF450 | GCF360, **GCF450**, GCF360 | Orfamide A* |
| CCMSLIB00004679299 | GCF450 | GCF360, **GCF450**, GCF360 | Orfamide B* |
| CCMSLIB00004679300 | GCF450 | GCF360, **GCF450**, GCF360 | Orfamide C* |
| CCMSLIB00000001573 | GCF468 | **GCF468**, GCF468, GCF235 | Barbamide |
| CCMSLIB00000001575 | GCF468 | **GCF468**, GCF468, GCF235 | Barbamide |
| CCMSLIB00000001706 | GCF471 | **GCF471**, GCF498, GCF550 | Jamaicamide A |
| CCMSLIB00000001708 | GCF471 | **GCF471**, GCF498, GCF550 | Jamaicamide C |
| CCMSLIB00000579285 | GCF476 | **GCF476**, GCF219, GCF235 | Albicidin |
| CCMSLIB00004681475 | GCF476 | **GCF476**, GCF219, GCF235 | Propionyl-albicidin |
| CCMSLIB00004681481 | GCF476 | **GCF476**, GCF219, GCF235 | Beta-methoxy-albicidin |
| CCMSLIB00004681486 | GCF476 | **GCF476**, GCF219, GCF235 | Carbamoyl-β-methoxy-albicidin |
| CCMSLIB00004681487 | GCF476 | **GCF476**, GCF219, GCF235 | Carbamoyl-β-methoxy-asn-albicidin |
| CCMSLIB00000840594 | GCF488 | GCF740, GCF740, GCF739 | Nevaltophin D |
| CCMSLIB00005724004 | GCF498 | GCF471, **GCF498**, GCF550 | Cryptomaldamide |

This kind of multi-omics analysis allows to dereplicate more metabolites than genomics or metabolomics separately and it helps researchers to focus on novel structures that can also present novel bioactivities.

We expect that our approach will improve with a larger training set and with further improvement of the features in the BGC and MS/MS fingerprints (e.g., by adding BGC regulation and predicted bioactivity). We confirmed that all 13 correct GCF predictions reported here were found in the original producer of the identified metabolites, they matched the reported masses and most of these BGC-metabolite links were reported at the PoDP as validated via knockouts, heterologous expression, and/or isolation and NMR structure elucidation. NPLinker (14), another recently published paired-omics tool, also uses validated links from the PoDP to assess linking precision scores. With 50 known GCF-MS/MS links that were present in the 1,040 samples with paired data (some metabolites have multiple MS/MS spectra), the annotation rate was reasonably good (around 28%, 14 out of 50 links were retained after the Jaccard co-occurrence threshold, a threshold to keep only the metabolites that are found among the same samples that contain the candidate BGCs). Table S1 provides an overview of the various numbers of genomes, metabolomes, BGCs, metabolites, etc.

### Dereplicating a new analogous MS/MS spectrum (with a library hit from GNPS but not found at the PoDP database) to a known BGC and accurately predicting its full *in silico* structure

In addition to dereplicating known metabolites (from the GNPS and/or the PoDP database), NPOmix can also dereplicate new putative analogs (orfamides marked by asterisks in Table 1). For example, the BGC for the metabolite orfamide C (genes 1-6 in Fig. S3, MIBiG ID BGC0000399) was automatically connected by our KNN approach (both using only similarity as a training/testing feature or by adding the biosynthetic class) to a GNPS metabolite labeled “putative orfamide C” (CCMSLIB00004679300). This MS/MS spectrum was also found in the same strain where the BGC was first identified (*Pseudomonas protegens* Pf-5, Genbank ID GCA_000012265)(27). The nine amino acid (AA) predictions for this BGC, based on the specificity of adenylation domains, match the structure for orfamide C in the correct order: leu, asp, thr, ile, leu, ser, leu, leu, and ser. AntiSMASH was not able to predict the tenth and last AA in the biosynthetic series, although, the mass difference between the partial structure and the experimentally observed *m/z* pointed that the last AA was indeed valine. The matching between the predicted structures (via dereplication versus *de novo* prediction using the NPOmix link) confirmed the multi-omics enabled dereplication of this “putative orfamide C” (using *k* = 3, BGC predictions and predicted metabolite structure are represented in Fig. S3), annotating this metabolite without the need for isolation. We would like to stress that this version of the KNN GCF predictions did not use structures/substructures for linking MS/MS spectra to BGCs; hence, as demonstrated in Fig. S3, substructure predictions can be an extra dimension for selecting links that are true positives over false positives.

### Connecting a cryptic/new MS/MS spectrum (not present in the GNPS database) to a known BGC (validated link from the PoDP database)

We used a combination of MS/MS fingerprints (notebook 2 from https://github.com/tiagolbiotech/NPOmix), BGC fingerprints (notebook 3), Mzmine (28), Dereplicator+ (29) to link and dereplicate brasilicardin A. After selecting 300 MS/MS spectra from the 16 most diverse genomes in the dataset (diversity estimated using GCF presence/absence) of 1,040 samples, Dereplicator+ had three *in silico* metabolomics predictions and one of them was the unique tricyclic glycosylated terpene brasilicardin A. The observed *m/z* matches the value previously reported in the literature)(30), identifying an MS/MS spectrum that is currently absent from the GNPS database. NPOmix connected the MS/MS spectrum (predicted to be brasilicardin A by Dereplicator+; please note that this information was not used during the NPOmix training) with the correct BGC (brasilicardin A MIBiG ID BGC0000632 from the strain *Nocardia terpenica* IFM 0406, GenBank ID GCA_001625105)(31), highlighting how NPOmix can connect cryptic metabolites without library matches (absent from MS/MS databases like GNPS) to their corresponding BGCs. Predicted fragmentation (Fig. S4) strongly suggested that the query MS/MS spectrum is corresponding to the planar structure of brasilicardin A (all deltas between fragmented exact *m/z* and observed *m/z* were extremely low, below 0.001, more information in Dataset S1, sheet four). Thus, NPOmix can provide links between cryptic MS/MS spectra and known/cryptic BGCs from the most diverse strains and potentially new BGCs that can be explored experimentally (e.g., BGC knock-out, heterologous expression, or isolation and NMR structure elucidation), especially if coupled to SMART-NMR analysis (32) to confirm their novelty. We observed that this compound was not in GNPS but it was already validated (via “knockouts, heterologous expression, or other gene cluster manipulation”, information at the dataset 1b0dccac-5212-4dfd-a9f2-6fa953ab16bd.5) and deposited in the PoDP database (same *m/z*, LC-MS/MS metabolome and retention time).

### Comparing the three different kinds of structural features for linking metabolites to BGCs using NPOmix

In order to increase the precision of our NPOmix algorithm, we added to the BGC and MS/MS similarity fingerprints the presence/absence of biosynthetic classes (polyketide synthases, nonribosomal peptide-synthetases, terpenes, siderophores, ribosomally synthesized and post-translationally modified peptides, phosphonates, oligosaccharides, phenolic metabolites, others/unknowns, other minor classes, and combinations of more than one class) and substructures (tyrosine, proline, amine, malonyl-CoA, and so on - all 85 predicted substructures can be found at Dataset S1, sheet five). A comparison of the various models is provided in Figure 3, with Fig. 3A without using the co-occurrence threshold, and Fig. 3B with use of the co-occurrence threshold.

**Fig. 3.**
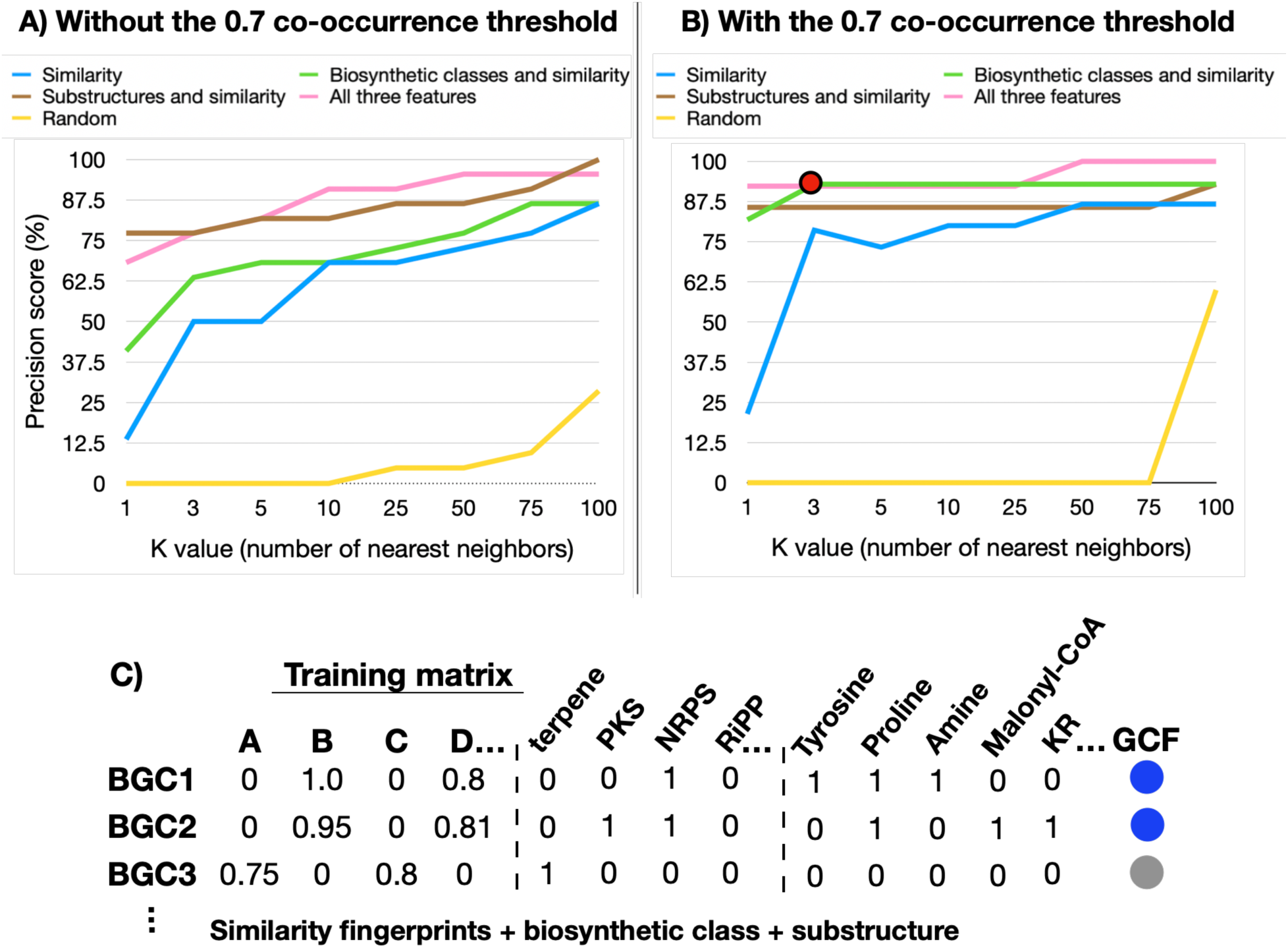
A) Precision curves without the Jaccard co-occurrence threshold for different features: similarity only (blue), biosynthetic class and similarity (green), substructure predictions and similarity (brown), and using both biosynthetic class, substructure predictions, and similarity (pink). B) Precision curves for the same cases using the co-occurrence threshold. Random (yellow) was calculated by using similarity only and shuffling the testing columns. The red dot indicates the best precision of 92.9% (using biosynthetic class and similarity as features, co-occurrence threshold, and *k*=3). *K* represents the number of nearest neighbors. C) we illustrate the modifications made in the training matrix with the addition of biosynthetic class and substructure prediction. For this work, we used 1,040 samples, therefore, 1,040 similarity columns plus 12 biosynthetic classes and 82 predicted substructures.

The comparison of the different features for running NPOmix is exhibited in Fig. 3, using the same set of samples previously described (1,040 paired samples and 22 validated BGC-metabolite links). As observed in the figure, the Jaccard co-occurrence threshold (a threshold to ensure that the query metabolite is connected to BGCs in the genome of the same microbial producer) substantially improved the precision scores, but it dropped the number of validated links from 22 to around 14 (these links are listed in Table 1). However, we obtained great scores even without threshold (Fig. 3A), for example, 81.8% for *k* = 5 using the three kinds of features (similarity, biosynthetic class, and substructure prediction) or just similarity and substructure features. Interestingly, we observed a slightly higher precision for *k* = 1 when only the substructures and similarity features are used in opposed to using all three kinds of features (without threshold); this is because the biosynthetic class for pyocyanines at MIBiG (used for manual annotation of the MS/MS spectrum) are annotated as “other” but antiSMASH automatically annotates its corresponding BGCs as “minor” and in some cases as both “minor” and “NRPS”, creating a mismatch between the biosynthetic classes. Additionally, we summarized KS domains and PKS substrates in only two columns (one for each) but we obtained the same precision scores. We elected the best precision score as 92.9% (red dot in Fig. 3B, score with co-occurrence threshold) because this very high value was obtained using only the similarity and the biosynthetic class features and three nearest neighbors, a good number of candidates for genome mining. Moreover, the use of just the similarity and biosynthetic class is very appealing because these kinds of features can be well predicted for metabolites with known or unknown metabolites, for example by using GNPS for similarity and CANOPUS and/or MolNetEnhancer for biosynthetic class predictions, as demonstrated by NPClassScore (33). It is much more challenging to predict substructures for unknown/cryptic metabolites, which is a topic of ongoing research by our group and others.

Fig. 3C, if a given BGC is a hybrid polyketide synthase (PKS) and nonribosomal peptide-synthetase (NRPS), it was annotated as 1 in the PKS and NRPS columns, and with a 0 in the remaining classes (additional columns). Analogously to the biosynthetic classes, each substructure prediction represented a new column in the BGC fingerprint or MS/MS fingerprints, and the columns were filled with 1 (if the substructure was present) and 0 (if the substructure was absent). For the MS/MS fingerprints in the testing set, we manually annotated these biosynthetic class features based on the known structures and MIBiG information. In cases where the structure is unknown, tools like CANOPUS (34) and MolNetEnhancer (35) can provide a similar biosynthetic class prediction relying on NPClassifier (36). For the substructure features in the MS/MS testing set, we used antiSMASH to annotate substructures for a BGC from the same GCF as the validated MIBiG BGC. In cases where the BGC is unknown, these predictions can be obtained: manually (by checking the known structure for common substructures); via unsupervised tools like MS2LDA (37), or; via supervised tools like MassQL (based on specific MS/MS fragments found in the spectra, manuscript in preparation) or CSI:FingerID/SIRIUS 4 (38). For example, we manually annotated the substructures predicted from the palmyramide A structure and we were able to correctly link the validated MS/MS spectrum with the corresponding palmyramide A BGC. This BGC is not yet published at MIBiG, but the biosynthetic gene cluster was previously annotated and reported in *Moorena producens* PAL (39), and in this study, the same genome, LC-MS/MS metabolome, and MS/MS spectral data were used to obtain this BGC-metabolite correlation.

### How well can we predict *de novo* structures from NPOmix links?

As observed by the four examples in Fig. 4, structural prediction can be very successful using multi-omics for peptidic metabolites without using library hits (the metabolite annotation such as orfamide C, and structure, was not provided in the training data for NPOmix); however, in some cases (like unusual amino acids in albicidin), it can be very difficult. We also showed that these *de novo* predictions are feasible to other classes, like the glycosylated terpene brasilicardin A (Fig. 4). Nonetheless, it is more challenging to predict the final structure (e.g., cyclization of the isoprene units in a diterpene) if compared to peptidic metabolites. As previously mentioned, we were able to correctly predict (*in silico* and *de novo*) the full structure for a new orfamide C MS/MS spectrum, including parts of its stereochemistry. This shows that NPOmix not only can dereplicate new MS/MS spectrum very efficiently, but it can also make very accurate predictions for cases where we did not use the library hits (in the MIBiG and/or GNPS databases). In other cases, we could only predict partial structure, as demonstrated by cryptomaldamide (85.7% correctly predicted), palmyramide A (66.6% correctly predicted), brasilicardin A (55.5%), and albicidin (29% correctly predicted). However, developments in the antiSMASH algorithm and the use of MassQL can almost certainly improve these predictions.

**Fig. 4.**
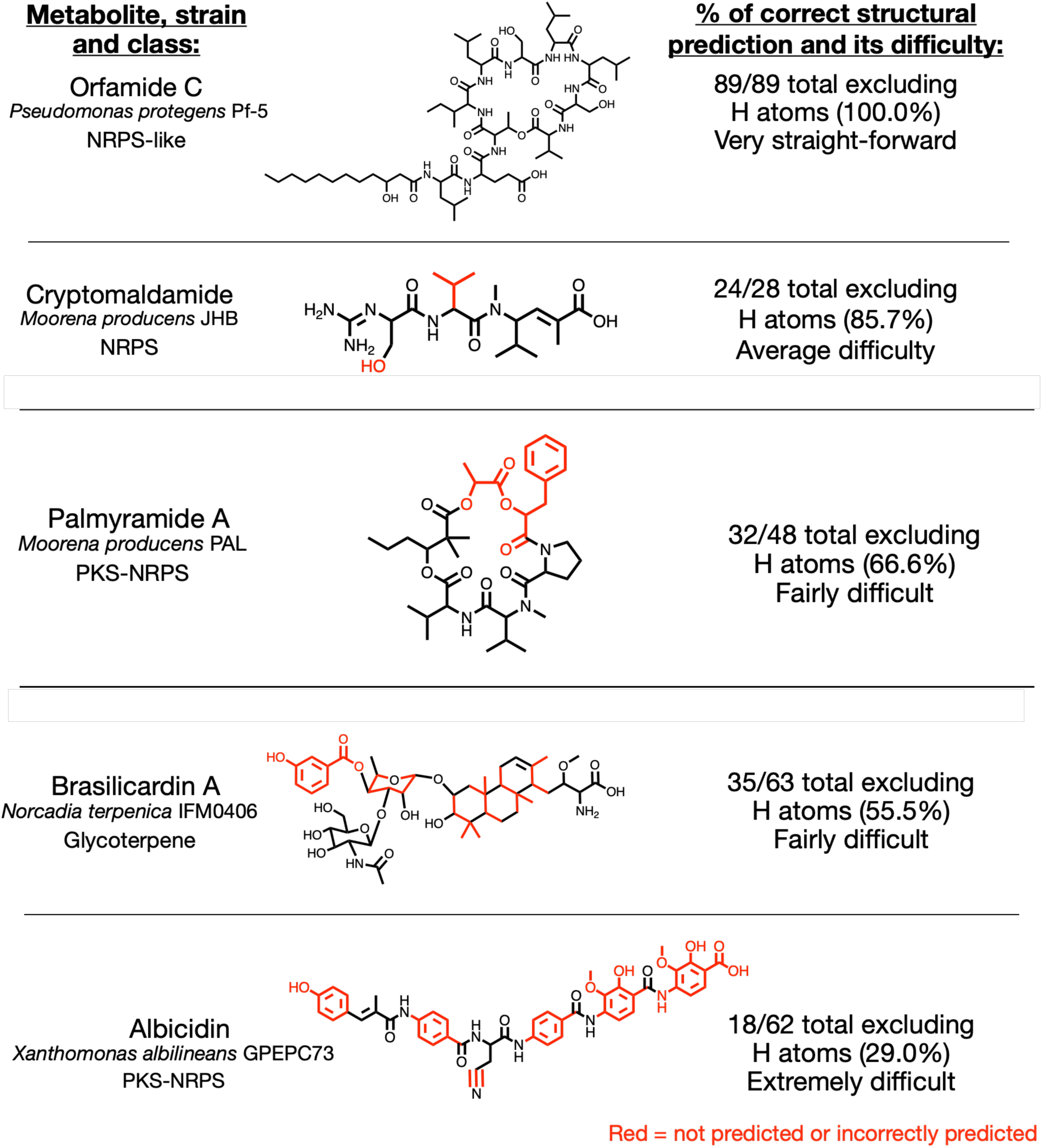
NPOmix automatically connected MS/MS spectra annotated with library hits to their peptidic MIBiG BGC (exemplified by orfamide C, cryptomaldamide, palmyramide A and albicidin), however, we did not use library hits for structure prediction. The figure illustrates the matches between the correctly predicted portion of the structures (atoms in black) versus the incorrectly predicted (substructures in red). We counted all atoms except for hydrogens (H). These predictions via antiSMASH used the BGC’s amino acids (AA) specificities and the PKS domains present in each investigated BGC. We would like to emphasize that these *in silico* predictions did not use the library annotations (obtained via dereplication or from the PoDP database) and, therefore, they are *de novo* predictions. In the structure of orfamide, only one AA (the last valine in the biosynthesis) out of 10 AA could not be predicted by the BGC annotation tool (antiSMASH), however, this valine residue was suggested by the *m/z* mass difference. In albicidin, unusual AAs are more difficult to predict than canonical AAs, leading to a lower number of correctly predicted atoms. BGC = biosynthetic gene cluster; AA = amino acid; H = hydrogen; PKS = polyketide synthase; NRPS = nonribosomal peptide-synthetase.

With MassQL (a tool data uses query language to find patterns in the MS2 data), specific mass spectral peak patterns that correspond to class-defining substructures or scaffolds can be mined from LC-MS/MS datasets (manuscript in preparation). Of note, we are mostly showing peptidic metabolites as examples because antiSMASH is currently better trained in predicting core structures from this biosynthetic class; nonetheless, we plan to expand this approach to other classes in the future, building on improvements in the genome mining field. Additionally, peptides are known for being bioactive (for example, orfamides and albicidins were reported as antifungal and the latter metabolite family was also reported as antibacterial), and they are likely the most abundant biosynthetic class in nature (accounting for 45% of the currently available BGCs at the antiSMASH database that contains over 82,000 BGCs)(40). Therefore, NPOmix links can already be used to explore environmental samples and make interesting *in silico* predictions about the structure and bioactivity of naturally occurring peptides.

### Comparing the performance of NPOmix with other published tools

We used the same previously described dataset of 1,040 paired samples that includes 22 validated BGCs-metabolites links to compare precision scores (same links used in the calculation of the precision scores in Fig. 3) to other 8 multi-omics (tools and their websites are listed in Table S3). For the tools that do not have a training step, we selected from the 1,040 paired samples only the ones that contained at least one of the 22 BGCs-metabolites validated links, a total of 9 paired samples. We compared NPOmix to NRPminer (12) and NPLinker (14), the other currently available tools that create links between MS/MS data and BGCs. NPLinker includes the correlation scores first developed in the Pattern-based Genome Mining publications (7, 8) and derivatives thereof. GNP (13) and NRPquest (11) seem to be discontinued (see Table S3 for further details).

MetaMiner (9) and DeepRiPP (10) could not be used in the comparison because, unfortunately, there are no ribosomally synthesized and post-translationally modified peptides (RiPPs) among the 22 links in our validation set. As far as we can tell, there are only two validated RiPPs in the entire PoDP database: capistruin C and polytheonamide A-B. However, neither of the two were available for our validation test (more details in the Supplementary Information, Results section). Under these circumstances, we did not test RiPP BGC-metabolite links, and we could not measure the performance of MetaMiner and DeepRiPP for this validation set, however, we tested both for the full metabolomes and genomes of these 9 validation strains. In this test, MetaMiner annotated zero matches, despite the presence of 49 uncharacterized RiPP BGCs in the 9 genomes. DeepRiPP “failed to load results”, even though the input data was correctly processed by other tools. On top of testing tools that connect mass spectrometry fragmentation data to BGCs, we also tested Nerpa (16) and GARLIC (17) which can connect structures (SMILES strings) to BGCs.

Fig. 5 is a histogram illustrating the performance of these tools, including the two best versions out of eight versions of NPOmix displayed in Fig. S5 (four with and four without co-occurrence threshold). Fig. S6 exemplifies how fragmented BGCs can lead to scores below the threshold. More details on the Fig. S5 and S6 in the Supplementary Information, Results section. We note that we did not observe differences in NPLinker results using the standardized Metcalf score, the Rosetta score or both together. We selected top-1 accuracy to represent the precision scores since some tools only provide a single BGC candidate per metabolite tested. In theory, NRPminer and Nerpa should be able to test four links that are nonribosomal peptides (three orfamides and nevaltophin D) plus eleven additional hybrids between nonribosomal peptides and polyketides (two barbamides, five albicidins, two jamaicamides, cryptomaldamide, and antimycin). However, NRPminer could only generate predictions for four of these peptidic metabolites in the validated set (one incorrect and three correct links) and Nerpa only predicted three metabolites that were correctly linked to their BGCs. GARLIC can test all but four links (three pyocyanines and brasilicardin A) since it should work for nonribosomal peptides, polyketides, and hybrids. We were unable to compile GRAPE (tool required to use GARLIC) and we found an unanswered issue on their GitHub page with that same question, as well as another issue from 2016 that is left unanswered, thus it seems that this tool is also no longer supported or discontinued.

**Fig. 5.**
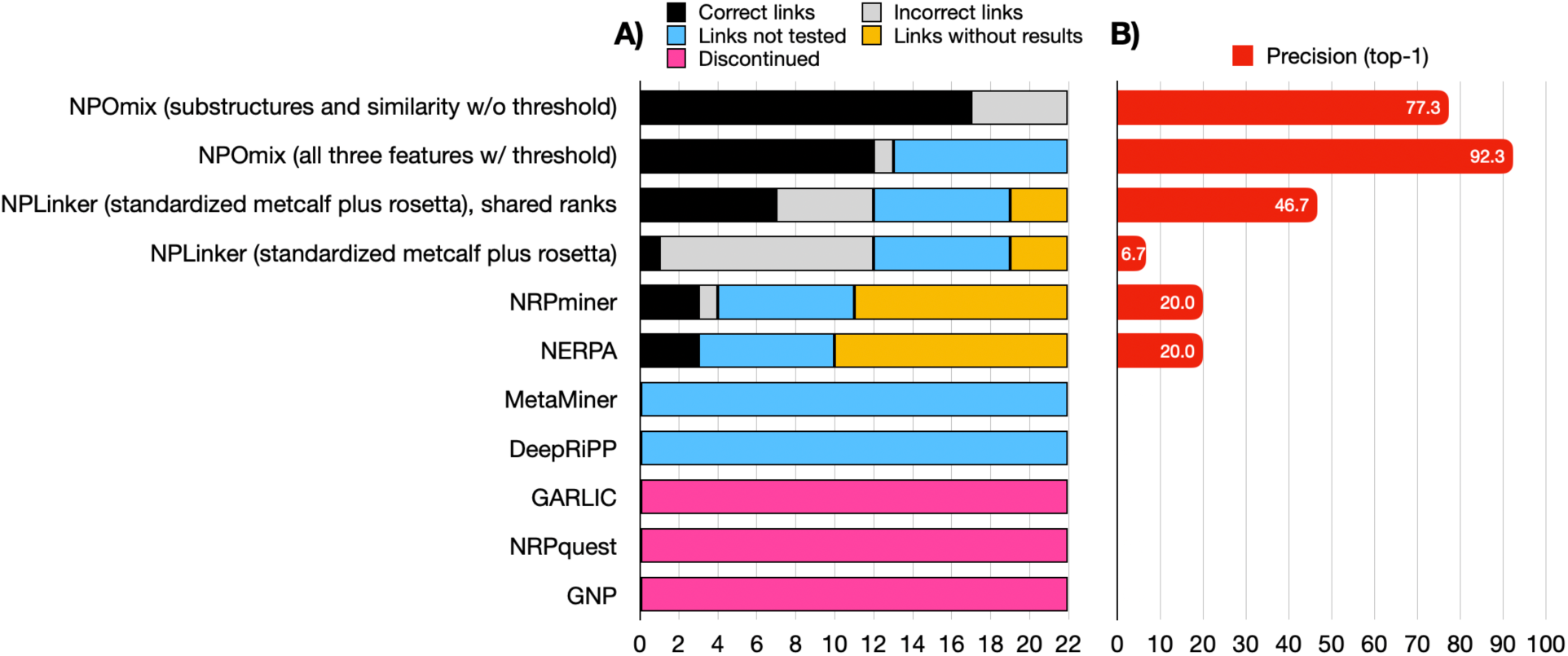
A) Histogram displaying the number of correct links (true positives, in black), incorrect links (false positives, in grey), links not tested (due to limitations of the tool or threshold selected, in blue), and links without results (in yellow) for different currently available multi-omics tools. NPOmix, NPLinker, and NRPminer can link MS/MS spectra (known or cryptic) to BGCs; GARLIC and Nerpa can link known structures to BGCs. NPOmix has different versions that yield different precision scores and so does NPLinker, however, in this case the scores were the same for the different versions. NPLinker has regular ranks and shared ranks (for example, from the second to the tenth position, if all scores were equal, then all of them would be tied for second). Precision is the ratio between true positive links overall tested positives (true, false, and links without result), and this metric was calculated using only the best candidate (top-1) from the tools’ outputs. NPOmix (all three features with co-occurrence threshold) had 9 links below the threshold and these links were not tested. NRPminer and Nerpa can only test peptidic metabolites. GARLIC and GNP were discontinued (in pink), and they could not test any of the links. NRPquest was replaced by NRPminer (hence, also discontinued, in pink). There were no known RiPP metabolites in the validation set for MetaMiner and DeepRiPP to test, however, these tools did not generate results for the 49 unknown RiPP BGCs found in the genomes from the validation set. W/ = with; W/o = without; RiPP = ribosomally synthesized and post-translationally modified peptides.

NPLinker is systematic (it does not depend on the BGC/metabolite class) and, just like NPOmix without threshold, it can test all links. For all tools, we used standard settings (commands found in the notebook “NPOmix_SI-installation_and_running”, GitHub repository https://github.com/tiagolbiotech/NPOmix); and we note that for these analyses we did not attempt to optimize settings and scores of the various tools but used their (recommended) default settings. We also note that the use of a relatively small number of strains for the NPLinker benchmarking purposes (9 strains), it is likely that the correct links are part of a longer list of possible matches based on their co-occurrence score and that several links have the same score. Hence, the correct match may not be in the top-1 rank if we do not consider a shared rank (for example, from the second to the tenth position, if all scores were equal, then all of them would be tied for second using their shared rank). The use of (many) more strains is likely to improve some rankings as the co-occurrence score is more likely to differentiate between different links. Both precisions for top-1, and top-1 shared ranks are provided in Fig. 5. The complete results list with regular ranking and shared ranking for NPLinker can be found at Dataset S1, sheet seven.

As illustrated in Fig. 5, the maximum number of correct links corresponded to using NPOmix with substructure and similarity scores without threshold, but the highest precision (92.3%) could be found using NPOmix with all three features (similarity, substructures, and biosynthetic classes) with threshold. This top precision from NPOmix in our benchmarking experimentation was about 46% more precise than the second-best tool tested, NPLinker with shared rank (the precision without shared rank was about 40% lower), and we were able to correctly predict about 2.5 times more BGCs-metabolites links than NPLinker. As aforementioned, five tools could not be tested because they were: discontinued like GNP and NRPquest; generated no results like MetaMiner; had unclear instructions like GARLIC; or could not read the same input that other tools were able to process normally like DeepRiPP. We concluded from this figure that NPOmix not only can predict more links (Fig. 5A) than most of the other tools but also the ratio of correct links is significantly higher than other methodologies (Fig. 5B). Of note, the prediction power of machine learning tools like NPOmix and NPLinker can be even higher with more top candidates (for this comparison we only used top-1) and better features, such as demonstrated by the implementation of biosynthetic class with NPClassScore (38) to improve NPLinker links and the use of extra features other than similarity by NPOmix. Machine learning tools can also improve with a bigger training set. In the future, we will compare NPOmix using the biosynthetic class feature with NPClassScore, since it would require standardizing and improving the current class predictions (using CANOPUS and MolNetEnhancer) for unknown/cryptic MS/MS spectra.

## Discussion

In this work, we demonstrated the use of machine learning (a new a K-Nearest Neighbors algorithm named Natural Products Mixed Omics, NPOmix) and genome mining for processing several thousand LC-MS/MS files and about a thousand genomes to connect MS/MS spectra to GCFs. Our approach can systematically connect MS/MS spectra from known metabolites (links validated experimentally), spectra from metabolites analogous to known (links with GNPS library matches, exemplified by orfamide C), and spectra from cryptic metabolites (links without GNPS library matches and therefore absent from the MS/MS database, as exemplified by brasilicardin A). The advantage of using paired data is that the genomic information represents the full metabolic potential of an organism, and hence, we can prioritize the discovery of the most diverse naturally occurring BGCs via genome mining. In other words, novelty predictions using BGC-metabolite links are likely more comprehensive than just using metabolomics and a more accurate picture of the actual chemical diversity in nature. Additionally, the use of genetic information can help in the structure elucidation and prediction of bioactivity (18), highlighting the advantage of using the BGC information in the drug discovery process. Moreover, predicting linked MS/MS spectra for a promising BGC can facilitate their heterologous expression as the expression can be difficult if the target metabolite is not known. We also show how cryptic MS/MS spectra (absent from MS/MS databases like GNPS) can be annotated using NPOmix, MZmine (28), and Dereplicator+ (29), allowing expansion of the current MS/MS databases.

NPOmix focus was on finding BGCs for metabolites, however, NPOmix could also be used to go vice-versa. Also, despite this manuscript being centered on how paired data can make better *in silico* predictions than just metabolomics, this same manuscript could also be focused on how paired data is better than just genomics and the benefits would be very similar. We believe this strategy aims to be a pipeline that better uses the wealth of available data; and therefore, it could maximize the chances of finding new drug-like metabolites.

We observed very good precision scores of top-1 = 81.8% and top-10 = 92.9% (both with randomness equal to 0, using the similarity and biosynthetic class features and using the Jaccard co-occurrence threshold). Additionally, we also obtained a good precision score of 77.3% even without the co-occurrence threshold (for *k* = 3 and all three kinds of features including similarity, biosynthetic class, and substructure predictions). We observed an annotation rate of around 28%, as 14 out of 50 MS/MS validated spectra that were retained after the co-occurrence threshold. The annotation rate was even higher without the use of the co-occurrence threshold, 44% (22 out of 50 dereplicated MS/MS validated spectra). Table S1 details the number of genomes, metabolomes, BGCs, GCFs and metabolites used in this study. More details on these 22 BGC-MS/MS links can be found in the Supporting Information (Discussion section) and Fig. S7.

In fact, compared to the other eight multi-omics tools, NPOmix seems to be the only available tool that is i) systematic (suitable to many classes of BGCs), ii) highly efficient (about 46% higher score in Fig. 5B than other approaches), iii) high-throughput (we processed around a thousand genomes and 3x more LC-MS/MS files), and iv) scalable (requiring low processing power, we used 8MB of RAM and 8 cores). We note that NPLinker is also systematic, high throughput, and scalable, however, its precision score during benchmarking was 46% lower than NPOmix, in part due to the relatively low number of strains used during NPLinker benchmarking. NPOmix was able to correctly predict about 2.5 times more BGCs-metabolites links than other tools. The combination of multiple multi-omics tools (and other genomics and metabolomics tools) can be a potent genome mining approach that would enable the visualization of many kinds of predictions (similarity scores for the links, biosynthetic classes, complete/partial structures, taxa, and bioactivity) in a single “omics network” (work in progress), prioritizing new true positives.

Of note, new links between metabolites with predicted structure and known BGCs may not mean that the metabolite fully corresponds to the predicted structure because mass spectrometry tends to not distinguish isomers and small changes in BGCs can yield substantial changes in the final structure. Therefore, NPOmix (a correlation-based approach) does not have the resolution to distinguish between isomers and regioselectivity of the various homologs. We would like to emphasize that new links (not experimentally validated links that are, for example, reported in the PoDP database) require proper elucidation to confirm the metabolite’s planar (2D) and absolute (3D) chemical structure. Hence, in this study, 6 of the 11 BGC-metabolite links (family of analogs) used for validation were previously fully elucidated (via knockouts, heterologous expression, and/or isolation and NMR structure elucidation) as recorded in the PoDP database. We want to emphasize that despite the small size of the validation set (11 BGC-metabolite links plus analogs, a total of 22 MS/MS spectra), we were able to combine *in silico* tools like Mzmine, Dereplicator+, and NPOmix to create new links that can expand MS/MS mass spectral reference database as level bronze metabolites (new putatively annotated MS/MS spectra). Another aspect to mention is that the precision score may not be reproducible for every dataset, especially if the dataset is not too similar to the current training set, as they may present very different features; and therefore, NPOmix features might not perform as well on a given atypical dataset. Interestingly, some of our *in silico* (*de novo*) structure predictions (such as for orfamides) were very accurate and they would substantially facilitate NMR structure elucidation of the naturally occurring metabolite.

The use of complete genomes over MAGs and metagenomes is preferred to create a more “complete” training set; we predict that this would result in better precision than if the training set is populated with several fragmented BGCs. It is important to clarify that some organisms do not have BGCs (e.g., higher plants and animals) and therefore their metabolites cannot be linked to biosynthetic genes using NPOmix. Regarding lower plants, the annotation of BGCs is now a hot topic, however, there is a current lack of paired data for plants and the current NPOmix demonstration is done on the suitable and available data from the PoDP (almost exclusively from bacteria). On top of that, BGC detection is more challenging for plants (e.g., more scattered biosynthesis genes).

Another challenging type of samples are microbiomes, such as non-axenic cyanobacterial cultures like the example of the mix between *Moorea producens* JHB and its 15 detected heterotrophic genera plus viruses (41). Although, we successfully processed 40 cyanobacterial microbiomes and we linked five metabolites to their correct BGCs, demonstrating that this tool can also be used with microbiomes. A major limitation of using microbiomes for genome mining is that the quality of their metagenomes can hinder BGC assembly, leading to a smaller number of BGCs than expected and many of these BGCs can be fragmented. We plan to address microbiomes, plants, algae, and fungi samples more comprehensively in our future work (other future goals also highlighted in the Supplementary Information, Discussion section).

Finally, we would like to stress that all true positive BGC-MS/MS validated links reported here were found in a known producer of the metabolites, they matched the reported masses, and six of the 11 BGC-metabolite links used for validation (true and false positives, including brasilicardin A) were reported at the PoDP as validated via knockouts, heterologous expression, or isolation and/or NMR structure elucidation. In a similar way, NPLinker also used such validated links from the PoDP database to assess the impact of novel BGC-MS/MS linking scores and the complementary of them. Palmyramide A was also previously validated via isolation and NMR structure elucidation (42) (hence, making seven out of 12 validated family links), and it will be soon published at MIBiG and PoDP. The putative orfamides were not confirmed via structure elucidation or co-elution with standards (the new metabolite spectra were annotated via GNPS library matching) but the comparison to a reference from GNPS and BGC amino acid predictions strongly indicates that they share the same planar structure as the reference metabolite from GNPS. Our results highlight the importance of making genomics and metabolomics data publicly available with curated metadata, because more available paired data would enable better training of models, and therefore, better tools for the research community.

## Conclusions

We developed a promising tool to search for new specialized metabolites in paired omics data of natural extracts by using links between cryptic MS/MS and cryptic biosynthetic gene clusters (BGCs) but also more efficiently dereplicating known BGCs or known metabolites. This will facilitate the use of genome mining in drug discovery pipelines. For example, we illustrated that our tool could integrate multiple kinds of information about biosynthetic classes, family beta-diversity, dereplication, and the probability of being a correct link. These bits of information are very useful to find novel metabolites from nature (cultures or metagenomic samples. Thus, we believe that the current version of NPOmix will already facilitate the access to the so far under-explored bacterial biosynthetic potential, moving from bioprospecting about 3% (1) to bioprospecting a larger portion of this potential.

Of note, we developed NPOmix and correctly used NPLinker (14), CANOPUS (34), NRPminer (12), Nerpa (16), GNPS molecular networking (13), BiG-SCAPE (6), and MetaMiner (9); these tools processed data from three databases (MIBiG, PoDP, and GNPS)(13, 18, 43). Interestingly, the information from linked BGC-MS/MS data was able to provide good predictions for partial structures like cryptomaldamide and even complete planar structures like orfamide C (including hints of the AA stereochemistry that can aid in the elucidation of absolute structures). Many of the links were known antifungal and antibiotic metabolites, therefore, reproducing this analysis for new metabolites can be a very promising methodology for drug discovery (work also in progress, especially focusing on the discovery of new siderophores).

To facilitate the use of NPOmix, we are hosting workshops (in English and Portuguese) and we created video tutorials, both available at https://www.tfleao.com/npomix1). We will assist the NPOmix use so the genome mining community can benefit from its capabilities, especially via GitHub (https://github.com/tiagolbiotech/NPOmix_python for the python version or https://github.com/tiagolbiotech/NPOmix for the Jupyter notebook version).

We anticipate the use of this pipeline for many applications including (but not limited to): 1) studying siderophores and iron cycling under nutrient limitation; 2) studying humans, mammals, plants, marine invertebrates, and other microbiomes and their relationship with host health; 3) finding new bioactive metabolites for drug development; 4) better understanding metabolite-mediated cell function of a model organism like *E. coli* (important for heterologous expression); 5) learning how plants, corals and phytoplankton can be more resilient to global warming and other anthropogenic impacts. In conclusion, this tool is going to be very important for the community, and it has a range of implications for genomics, metabolomics, and other research fields.

## Code and Data Availability

The code (a collection of Jupyter notebooks) required to reproduce this work and to use the NPOmix tool for new samples can be found on the following GitHub repository page: https://github.com/tiagolbiotech/NPOmix. The repository also includes short video explanations on how the tool works and its importance for natural product discovery. We also wrapped the code in a few python scripts found here: https://github.com/tiagolbiotech/NPOmix_python. This page with the python version also includes a video explanation of how to prepare inputs and how to run NPOmix in python.

Additionally, we provide a webpage (https://www.tfleao.com/npomix1) for submitting a limited number of samples for free processing (maximum of 50 queries MS/MS fragmentation spectra) and this page also includes workshops on how to submit samples and interpret results. We are going to purchase a server and allow users to submit many more metabolites than just 50.

The (meta)genomes used to create the NPOmix training dataset for validation were downloaded from the Paired omics Data Platform (PoDP)(21) using notebook 1 from the GitHub repository (Jupyter notebook version). The paired experimental MS/MS files were downloaded using the ftp links (also from the PoDP) found in Dataset S1, sheet six. The testing set included MS/MS spectra from PoDP, spectra from the Global Natural Products Social Molecular Networking database (GNPS)(13), and also spectra also used in the NPLinker publication (33). If the potential users find the tool challenging to run, we have our contact information on the main webpage (link above) to submit samples and we expect that promising results will lead to fruitful collaborations.

## Funding and Acknowledgments

This research was supported by National Institutes of Health (NIH) Grants GM107550 (to P.C.D., W.H.G., and L.G.) and GM118815 (to W.H.G. and L.G.). This work was supported in part by a seed grant from the Center for Microbiome Innovation at the University of California San Diego. R.S. was supported by the São Paulo Research Foundation (Awards FAPESP 2017/18922– 2 and 2019/05026–4). J. J. J. v. d. H. and J.J.R.L. were supported by the Netherlands eScience Center (ASDI eScience grant ASDI.2017.030). J. J. J. v. d. H. was supported by the Netherlands eScience Center Open eScience Call (NLESC.OEC.2021.002). A.G. was supported by the System Biology Fellowship of Skoltech & PMI (Research Agreement number 8299). We would like to acknowledge Sandra Neubert, Bohan Ni, Dr. Kiefer Forsch, Dr. Christopher Leber, Dr. Chen Zhang, Dr. Andres Mauricio Caraballo-Rodriguez, Dr. Louis-Félix Nothias, Dr. Benjamin Naman, and Dr. Mitja Zdouc for the productive discussions about the topic. Additionally, we would like to acknowledge Dr. Andrew Ramsay for the help and advice on how to run NPLinker.

## Methods

See Supplementary Information Methods section for details on obtaining the data, assembly, BGC comparison, the NPOmix tool itself and other bioinformatic analyses.

## Supplementary Information

### Background

Besides the fact that our validation set is small (11 BGC-metabolite links plus analogs, a total of 22 MS/MS spectra), we do show that the combination of *in silico* tools like Mzmine (28), Dereplicator+ (29) and NPOmix can create new links that can expand MS/MS databases as level bronze metabolites (new putative MS/MS spectra). Whilst >2000 validated BGCs are described in the MIBiG database, a minority of them have associated MS/MS spectra, and a subset of these can be expected to be present in samples under study for which actual paired omics data is available, explaining why the validation set is small). Nevertheless, we were able to correctly connect 17 known metabolites (including analogs) to their validated BGCs using their similarity, biosynthetic class, and substructure features. We also exemplified with orfamide C (peptidic metabolite) how the information from these links can yield very accurate *in silico* and *de novo* planar structure prediction with a low delta *m/z* (error between predicted and observed *m/z*). We believe the same approach can be used for new/cryptic metabolites since the KNN algorithm does not know the metabolite labels (e.g., orfamide C) while making the BGCs-metabolites links. Moreover, we could partially predict the stereochemistry of orfamide C and other metabolites using genomic data.

Generally, finding novel metabolites for cryptic BGCs or even known BGCs (e.g., new analogs) is very useful to accelerate natural product discovery, however, connecting known metabolites to their biosynthetic gene clusters is also important. Newly linked BGCs for known metabolites can lead to the discovery of new enzymatic processes. For example, in the strain *Anabaena variabilis* ATCC 29413, a nonribosomal peptide-synthetase (NRPS) gene is responsible for the attachment of a serine residue to generate the final mycosporine-like amino acids (MAA) product. However, in the strain *Nostoc punctiforme* ATCC 29133, this same step is performed by an ATP-grasp ligase (44). This highlights that different microbes can generate the same metabolite through different convergent biosynthetic routes.

## Results

### Comparing the performance of NPOmix with other published tools

Capistruin C was “bioinformatically inferred through comparative analysis with another experimentally defined gene cluster”. Therefore, capistruin C was not validated in the reported strain at PoDP (*Burkholderia thailandensis* E264). The strain in which the BGC was validated is absent from the MassIVE database (*Burkholderia thailandensis* E444), meaning that we could not obtain the metabolome required for testing this strain. Moreover, we believe that this prediction “inferred through comparative analysis” does not need to be correct because the *B. thailandensis* E264 genome was included in the NPOmix analysis but the MIBiG BGC did not network with any other BGC in the full dataset (including the MIBiG BGC known for producing capistruin C), and we also checked the *B. thailandensis* E264 antiSMASH result for a capistruin C annotation but, again, none of its BGCs had the capistruin C annotation as the “most similar known cluster”. We were unable to find the MS/MS spectra for polytheonamide A and B, they are not published at GNPS.

As observed in Fig. S5A, the absolute number of correct BGC-metabolite links is bigger for NPOmix using the substructure and similarity features without the co-occurrence threshold compared to the same condition using the co-occurrence threshold, because some links are present in several samples and some BGCs can be silent or very fragmented. This typically creates mismatches between the fingerprints and drops the Jaccard index in the presence/absence of the fingerprints used for the co-occurrence threshold. However, some not so fragmented BGCs can be connected via BiG-SCAPE similarity to a reference BGC and then get grouped into the same family, see for example in Fig. S6. Additionally, in some samples, the MS/MS spectra might not be acquired due to the complexity of the sample and the nature of the data-dependent acquisition. Hence, in these cases, the co-occurrence threshold may remove links (illustrated as “links not tested” in Fig. S5A and Fig. 5) even if they are true positives (metabolite correctly linked to its BGC). For NPOmix using similarity only and using both similarity and biosynthetic class, the threshold only removed false positives. Despite the higher number of correct links without the threshold, the threshold increases the relative ratio of correct links, leading to higher precision scores if compared to the NPOmix versions without threshold (Fig. S5B).

## Discussion

We created a machine learning solution, a K-Nearest Neighbors algorithm named Natural Products Mixed Omics (NPOmix) approach, to connect natural products observed by untargeted mass spectrometry to their corresponding biosynthetic gene clusters (BGCs). We showed that a large dataset, deriving from heterogeneous sources such as the ones currently available in the Paired Omics Data Platform (PoDP), can create good fingerprints and can thus successfully connect known metabolites to their corresponding BGCs, such as albicidin and its analogs to a BGC in *Xanthomonas albilineans* GPE PC73 (GenBank ID GCA_000087965.1), orfamides A-C to a BGC in *Pseudomonas protegens* Pf-5 (GCA_000012265), and cryptomaldamide and jamaicamide A and C to BGCs in *Moorena producens* JHB (GCA_001854205). All three of these strains were the original producers of these metabolites. In Fig. S3, we illustrated how the BGC predictions (such as predicted moieties) can help to prioritize true links over false positives via comparing predicted structures between a given MS/MS spectrum and its BGC candidates (the matching was done using only similarity and biosynthetic class as features for the NPOmix algorithm).

We are developing an integrated pipeline for metabolite discovery using genomics, LC-MS/MS metabolomics, and other multi-omics tools. Additional future work will include the testing of other similarity metrics for networking and fingerprinting such as BiG-SLICE (45) for genomics and Spec2Vec (46) and MS2DeepScore (47) for the metabolomics. We will also look for synergy with correlation scores from NPLinker (14) to better annotate paired omics datasets. We intend to implement biosynthetic class and substructure predictions straight from the MS/MS fragmentation spectra using tools like SIRIUS 4 (38), MS2LDA (37), MolNetEnhancer (35), or CANOPUS (34), prioritizing candidates that have several substructures or predicted classes matching between BGCs and MS/MS spectra. The GNPS molecular family information could be used to select a consensus prediction among different MS/MS spectra from the same family. Enrichment of the current Paired Omics Data Platform dataset (we could now use 1,040 PoDP samples for NPOmix) with higher quality samples as well as more validated BGC-MS/MS links will further drive the development of tools such as NPOmix, and this will spark the discovery of more novel NPs.

We intend to add to this multi-omics approach: 1) BGC bioactivity (20); 2) MS/MS bioactivity, by creating a new machine learning tool to predict bioactivity straight from the MS/MS spectra, and; 3) MS/MS substructure predictions, by integrating tools like MS2LDA (37), CSI: FingerID /SIRIUS 4 (38) and MassQL (not published yet). NPClassScore (33) demonstrated that biosynthetic class can be predicted with a combination of CANOPUS and MolNetEnhancer, hence, the NPOmix users can already run the KNN version with similarity and biosynthetic class as features, a version that yielded a precision of 92.9% in the validation set. Accurate *in silico* structure and bioactivity predictions allow start using a more “modeling” way to investigate all sequenced environmental bacteria/microbiomes, reducing the need for multiple cell assays and significantly aiding structure elucidation.

NPOmix was able to correctly link nine out of 12 different known metabolites (18 out of 23 MS/MS spectra, including analogs) to their corresponding BGCs. These metabolites (and analogs) were barbamide, antimycin A1, pyocyanine, brasilicardin A, orfamide A-C, albicidins, jamaicamide A and C, cryptomaldamide and palmyramide A (the last one was tested separately from the validation set and it was not used to estimate precision scores in Fig. 3 and 5). These known metabolites were reported from 9 strains belonging to the phyla Proteobacteria (44.44%), Actinobacteria (33.33%), Cyanobacteria (22.22%), and from seven different genera (Fig. S7A), indicating taxonomic diversity in this bacterial validation dataset. Most of these links were reported to be active against fungi and bacteria (Fig. S7B). We can also see in Fig. S7B that these links encompass distinct biosynthetic classes like polyketide synthase (PKS), nonribosomal peptide-synthetase (NRPS), hybrid PKS-NRPS, hybrid terpene-oligosaccharide, and other; indicating that the tool is indeed systematic.

## Methods

### Obtaining paired data

The paired data used in this manuscript is available at the Paired omics Data Platform (PoDP) and we were able to obtain 36 out of 71 meta-datasets that were available at the time. We automatically downloaded the paired (meta)genomics-metabolomics data from the samples in the PoDP according to the code in notebook 1 at the GitHub repository described below.

### Metagenome assembly and annotation, BGC, and MS/MS similarity calculation

Metagenomic reads were assembled with SPAdes 3.15.2. (48). For BGC annotation, we used antiSMASH 5.0 1. (5) and for gene cluster networking we used BiG-SCAPE 1.0 (similarity cutoff of 0.7)(6). BiG-SCAPE raw distance is measured via the domain sequence similarity (DSS) index, an index that calculates the Pfam domain copy number differences and sequence identity. For networking metabolites, we used GNPS classical molecular networking release 27 (similarity cutoff of 0.7). We did not use the full classical molecular networking capabilities in the NPOmix approach, as only the functions required to calculate a modified cosine score between a pair of MS/MS spectra were needed.

### Creating fingerprints

We developed python scripts and we combined them with scripts from sklearn (https://scikit-learn.org/stable/index.html) to create both BGC and MS/MS fingerprints and to run the KNN algorithm. A BGC fingerprint is created by pairwise BiG-SCAPE comparison between the queried BGC and all the BGCs found in the (meta)genomes in the training set, selecting the highest similarity scores for each (meta)genome. An MS/MS fingerprint (part of the testing set) is created by pairwise modified cosine comparison between the queried MS/MS and all the MS/MS present in the LC-MS/MS files paired with the genomes from the training set, also selecting only the highest similarity scores per set of experimental MS/MS spectra.

### Jupyter notebooks

All scripts used in this research can be found at this GitHub repository: https://github.com/tiagolbiotech/NPOmix. For instructions on installing and running the tools used in this publication, please see “NPOmix_SI-installation_and_running”. Notebook 1 can be used to download (meta)genomes and metagenome-assembled genomes (MAGs) that contain paired untargeted metabolomics (LC-MS/MS)(metabolomic files will also be downloaded by the notebook). We selected genomic samples that contained a valid Genome ID or BioSample ID, resulting in 732 genomes/MAGs. We also selected and assembled 1,034 metagenomes.

Notebook 2 can be used to process downloaded metabolomics files and a selected set of “.mgf” reference MS/MS spectra, creating a matrix containing the MS/MS fingerprints for the selected set of reference spectra (reference MS/MS spectra for the validation but for using the tool these reference spectra will be replaced by cryptic MS/MS spectra). If there were more than one LC-MS/MS file per genome (for example different media conditions or different chemical fractions), these files were merged into a single file representing these experimental MS/MS spectra. Notebook 3 can be used to process the antiSMASH results to create BGC fingerprints and use those to train the KNN algorithm. The MS/MS fingerprints are used to predict a/multiple GCF(s) for each tested reference MS/MS spectra found in the paired genomes-MS/MS data. We filtered the GCF-MS/MS links for cases in which the top GCF candidate had co-occurrence with a cutoff of 0.7 (GCF and MS/MS scores were present in the same set of samples (the cutoff is the same as the similarity cutoffs for consistency). Notebook 3 also performs cross-validation (dividing the data into 5 parts) and the average precision score for the cross-validation was 56.9%. Notebook 4 can be used to generate metadata such as the type of GCF or the count of BGCs per each genus in the database. Notebook 5 presents the code for genome mining that yielded the annotation of brasilicardin A (more details below). Notebook 6 expanded the similarity/absence fingerprints by including the biosynthetic class as new features, notebook 7 included the substructure predictions as new features and notebook 8 uses all three features (similarity, biosynthetic class, and substructures).

### Genome mining for new MS/MS spectra using Dereplicator+ and NPOmix

In order to use the NPOmix approach to find new NPs without any GNPS library matches (absent from the MS/MS database), we developed a pipeline combining NPOmix, MZmine (28), and Dereplicator+ (29). First, several strains can be selected using MZmine, here exemplified with 16 strains, based on their BGC beta-diversity scores. The Jaccard beta-diversity score metric of the similarity between a pair of strains was calculated as the intersection over the union of the detected gene cluster families. Using MZmine, we select peaks that were above a certain intensity threshold (we used base peak relative abundance of 1E6) to prioritize the chromatographic peaks that could reasonably be isolated for structure elucidation. In this example, we detected approximately 3,800 peaks with MS/MS spectra found in the analysis of the 16 most diverse strains. This MZmine list of peaks that have associated MS/MS data was filtered for a minimum precursor mass of *m/z* 500 to promote the presence of multiple moieties (substructures) in the predicted structures, generating 300 “.mgf” files. These mgf files were used by NPOmix to predict the GCFs/BGCs for each of the 300 MS/MS spectra. We filtered for BGC-MS/MS links that the query MS/MS spectra existed in the same strains that the query BGCs were found and not across different strains, using the Jaccard index in the presence/absence of the fingerprints, essentially a pairwise analysis between the BGC fingerprint and the MS/MS fingerprint. This second filter narrowed down the number of mgf files to 72. These 72 mgf files were processed by Dereplicator+ for predicting structures for each MS/MS spectrum, leading to the annotation of brasilicardin A. Two other Dereplicator+ hits did not match the predicted GCFs. MZmine parameters were as follows: noise level of 1E6 for MS1 and 1E3 for MS/MS, minimum group size in the number of scans of 4, group intensity threshold of 1E6, the minimum highest intensity of 3E6, *m/z* tolerance of 10 ppm, retention time tolerance of 0.2, weight for *m/z* of 75%, and weight for a retention time of 25%.

### Expanding BGC and MS/MS fingerprints using biosynthetic classes

In notebook 6, the BGC classes were annotated and included in the BGC fingerprints. To accomplish this, all of the antiSMASH annotations for a given BGC were added to the presence of all predicted classes. Each class represented a new column in the fingerprints and the columns were filled with 1 (if the class was present) and 0 (if the class was absent). We observed the following classes in our dataset: polyketide synthases, nonribosomal peptide-synthetases, terpenes, siderophores, ribosomally synthesized and post-translationally modified peptides, phosphonates, oligosaccharides, phenolic metabolites, others/unknowns, other minor classes (aminoglycoside/aminocyclitol, beta-lactone, beta-lactam, butyrolactone, ectoine, furan, fused, indole and phenazine), and combinations of more than one class (like PKS-NRPS). We added in total 10 new presence/absence biosynthetic features. In the MS/MS fingerprint, for each one of the 22 validated MS/MS spectra, we annotated the presence/absence of the biosynthetic classes based on the known structures. These new fingerprints were used in the machine learning process, analogously to the notebook 3.

### Expanding BGC and MS/MS fingerprints using substructure prediction

The substructure predictions were also annotated using antiSMASH and they were implemented in the BGC fingerprints. Analogously to the biosynthetic classes, each substructure prediction represented a new column in the fingerprints and the columns were filled with 1 (if the class was present) and 0 (if the class was absent). We annotated 85 substructures (the substructures found in our samples are listed in Dataset S1, sheet five) and these substructures were added as 85 new features. We also tested summarizing and PKS substructures (malonyl-CoA, methylmalonyl-CoA, and so on) and all KR domains into two columns (one for each). Regarding the MS/MS fingerprints for each of the 22 validated MS/MS spectra, we annotated the columns based on the information from the known BGCs. For palmyramide A we manually annotated the substructures from its known structure. The new BGC and MS/MS fingerprints were used in the machine learning algorithm to calculate precision scores. Predicted structures were obtained by analyzing the substrate-specific from the antiSMASH BGCs and manually matching them to the MS/MS fragmentation spectra.

### Benchmarking multi-omics tools (comparing the performance of NPOmix with other tools)

We attempted to execute eight multi-omics tools for benchmarking NPOmix: MetaMiner (9), DeepRiPP (10), NRPquest (11), NRPminer (12), GNP (13) and NPLinker (14), Nerpa (16) and GARLIC (17). For the sake of computational resources, we only performed the benchmarking on the 9 strains that contained the 22 BGC-metabolite-structure links used for NPOmix validation and for calculating the NPOmix precision scores. We selected standard configurations for all these eight multi-omics tools and, unfortunately, we were unable to run DeepRiPP (could not process the FASTA files), NRPquest (discontinued), GNP (discontinued), and GARLIC (we were unable to compile its pre-requirement, GRAPE, and it seems that this tool was also discontinued). MetaMiner produced no results, despite the presence of 49 unknown RiPP BGCs in these 9 samples.

---

**Fig. S1.**
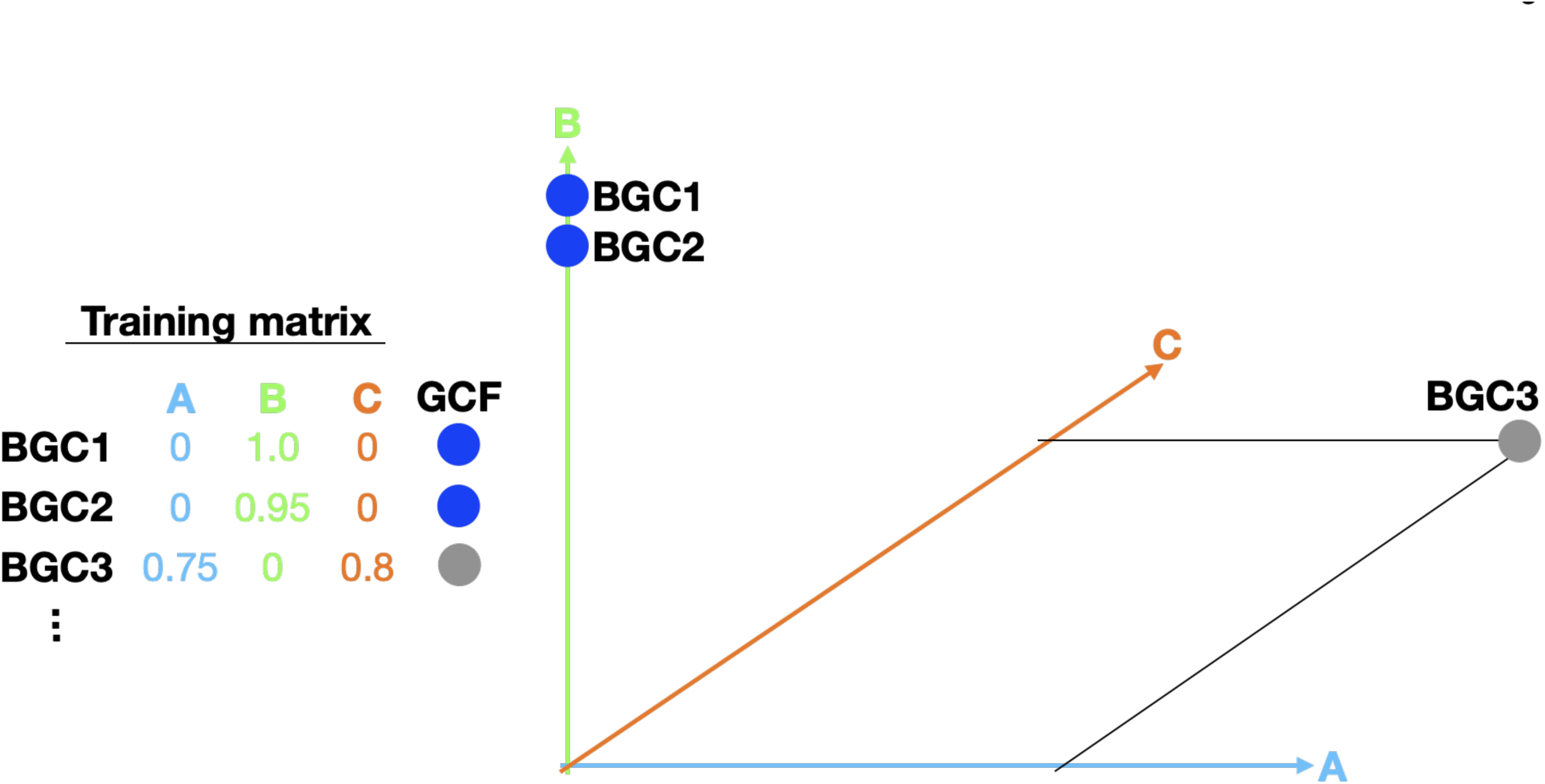
Representation of how BGCs can be plotted in the KNN (*k*-nearest neighbor) space by using the values in the training matrix, each column represents a genome in the training set and it also represents a dimension in the KNN space (1,040 genomes distributed in 1,040 columns). This example has three dimensions because it uses only three genomes; the actual training matrix used in this study had 1,040 genomes and therefore 1,040 dimensions. BGC = biosynthetic gene cluster; GCFs = gene cluster family.

**Fig. S2.**
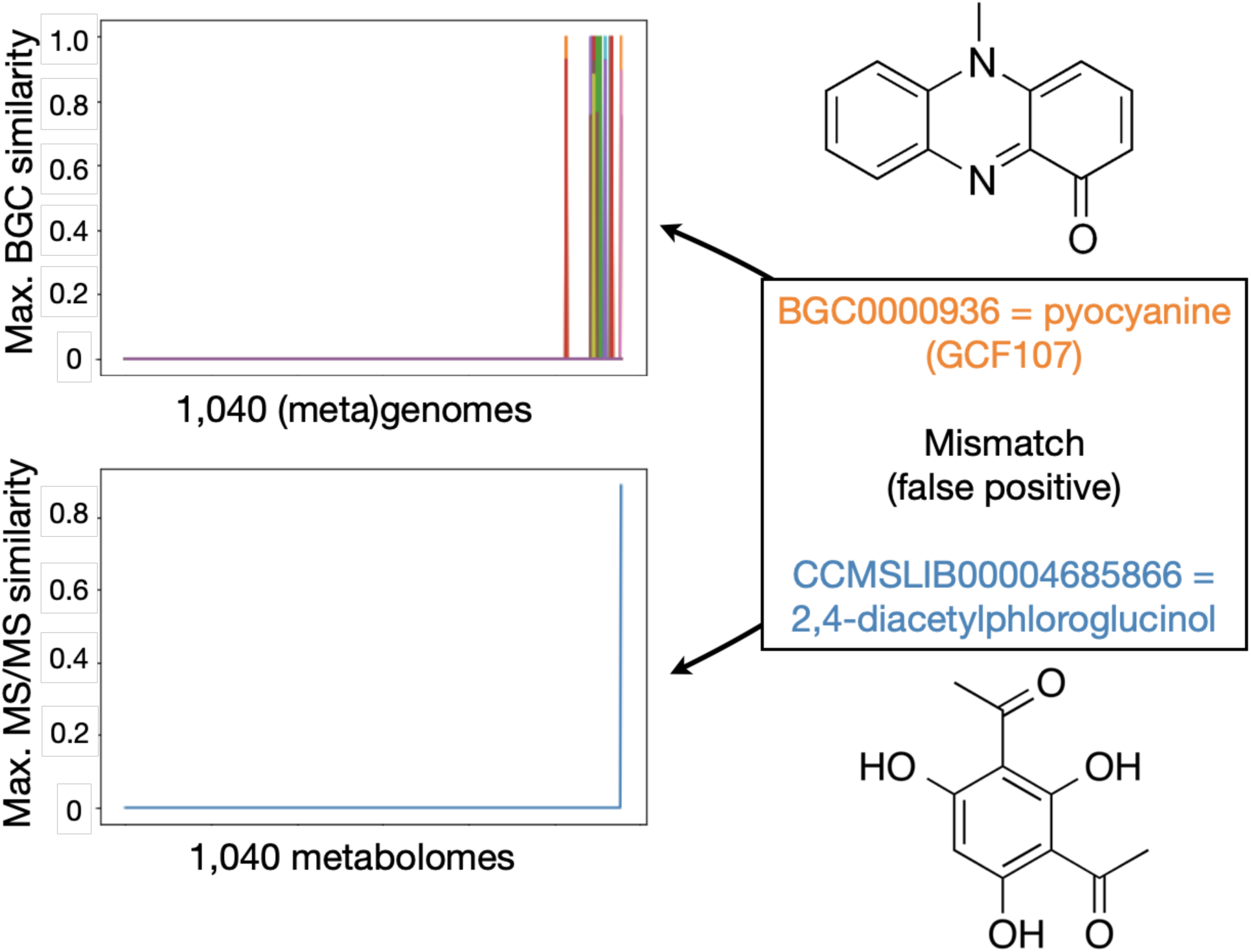
Representation of a mismatch linked by the KNN algorithm using *k* = 3. It is visually clear that the closest neighboring BGC fingerprints for pyocyanine do not properly match the MS/MS fingerprint from the metabolite 2,4-diacetylphloroglucinol, indicating that NPOmix suggested the wrong GCF for the 2,4-diacetylphloroglucinol MS/MS spectrum. BGC = biosynthetic gene cluster; GCFs = gene cluster family.

**Fig. S3.**
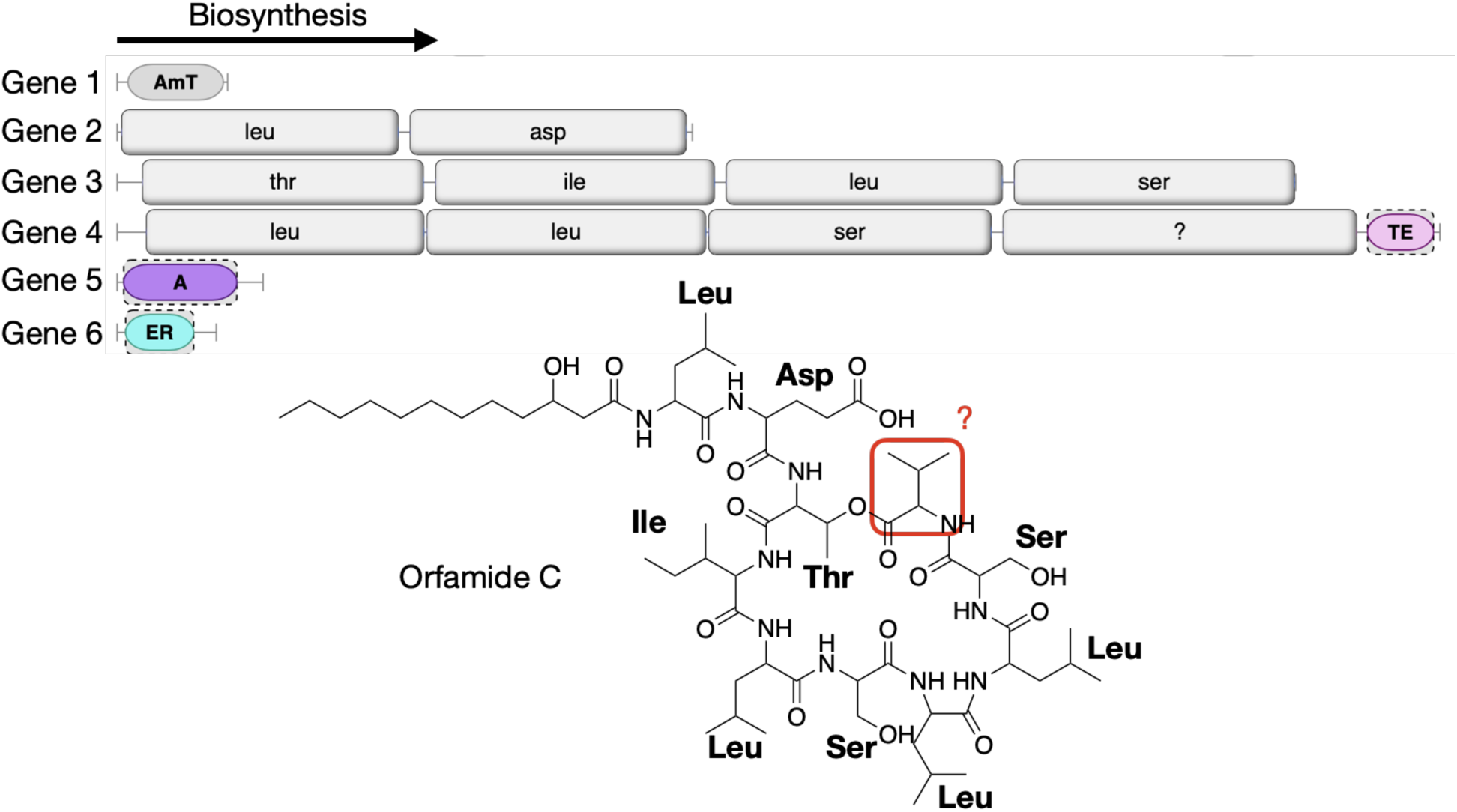
NPOmix automatically connected an MS/MS spectrum annotated as “putative orfamide C” to the MIBiG BGC annotated as orfamide C. The figure illustrates the matches between the BGC’s AA predictions (via antiSMASH) and the predicted metabolite structure (orfamide C, predicted via MS/MS spectral matching). Only one AA (valine, in red) out of 10 AA could not be predicted by the BGC annotation tool (antiSMASH), however, this valine residue was predicted by the MS/MS spectrum. BGC = biosynthetic gene cluster; AA = amino acid; AmT = aminotransferase; TE = thioesterase; A = adenylation domain; ER = enol reductase; “?” in the BGC represents that one AA could not be predicted by antiSMASH.

**Fig. S4.**
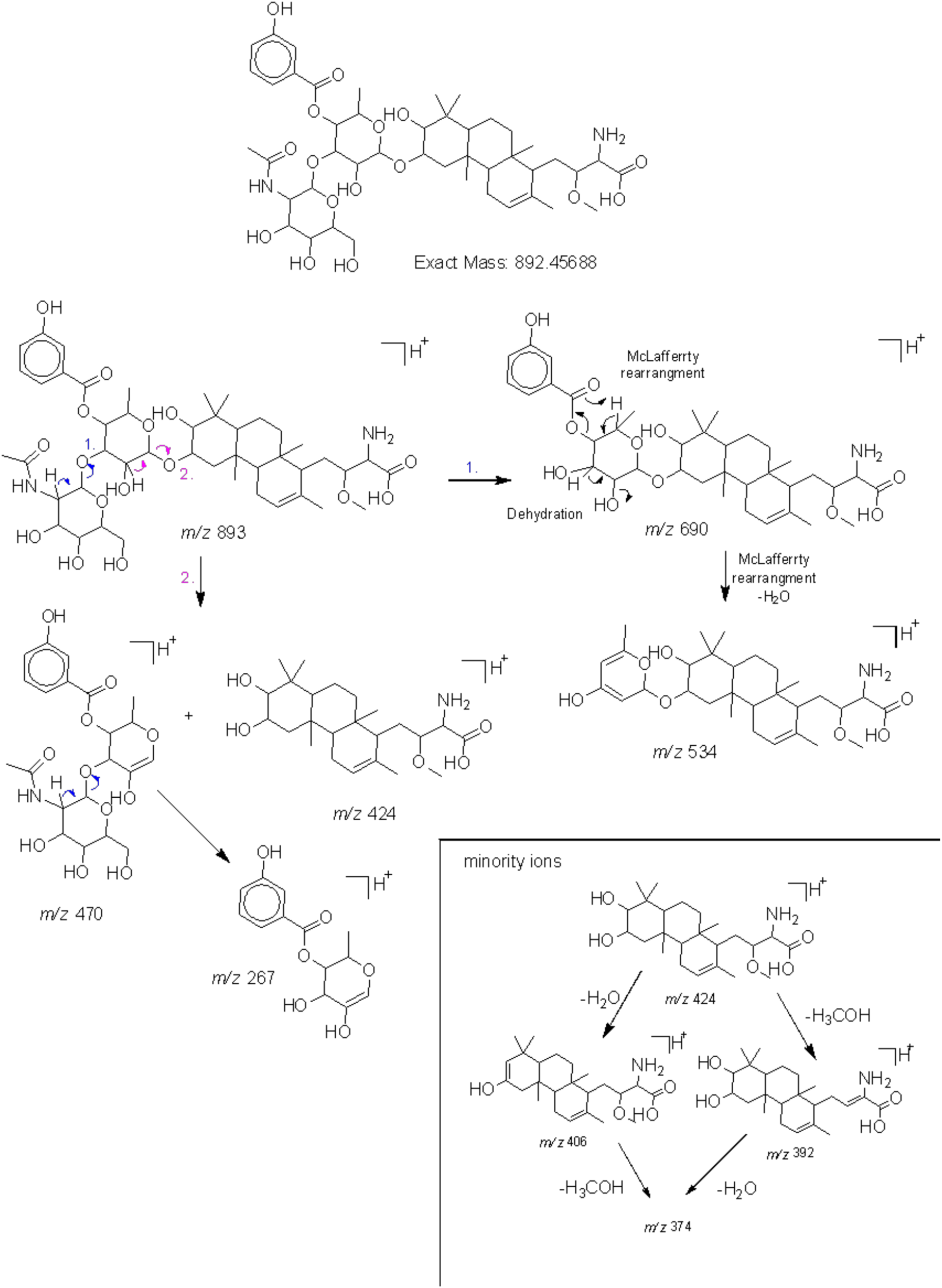
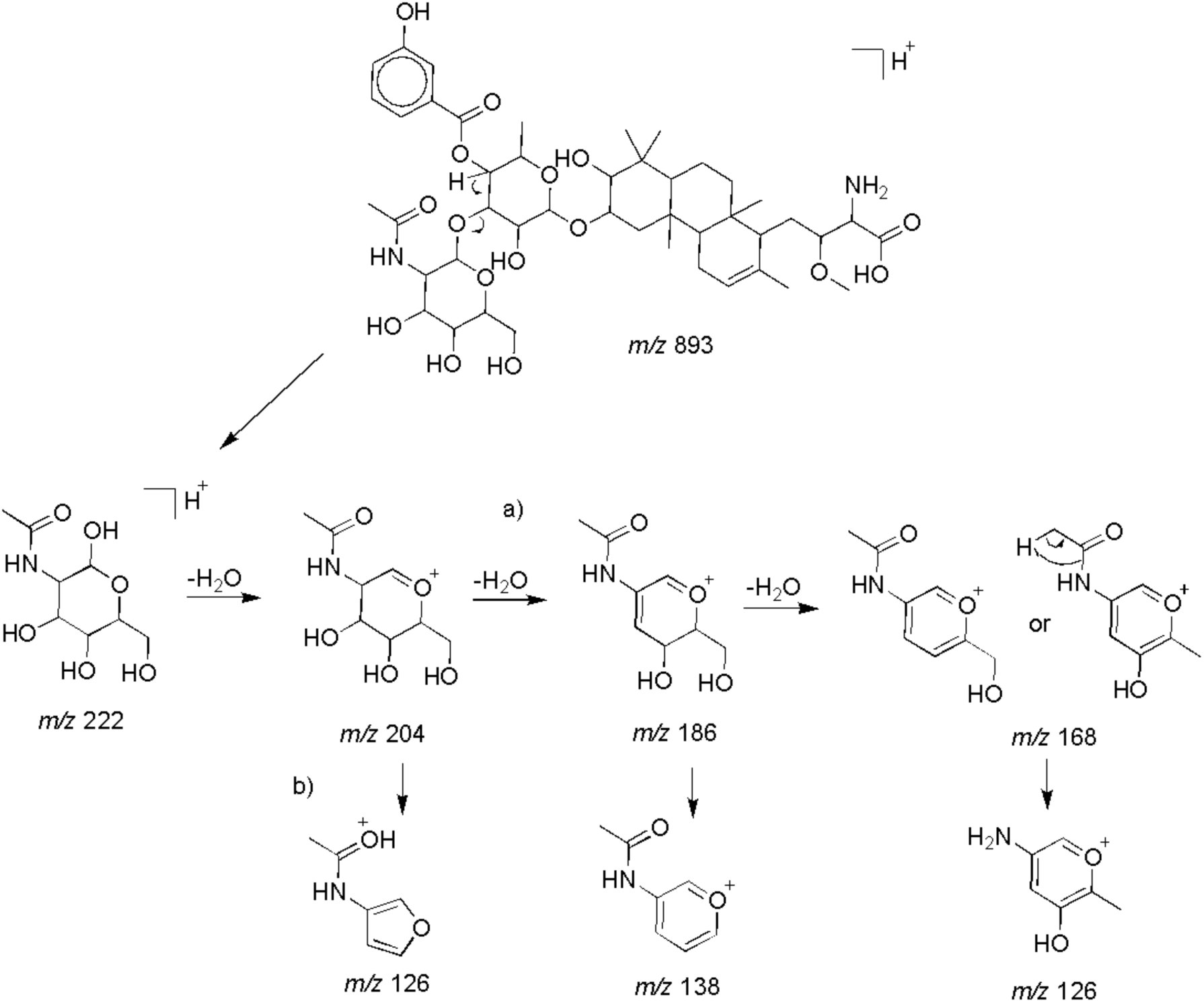
Proposed mechanism for the fragmentation of brasilicardin A by ESI mass spectrometry. The structure was proposed by NPOmix as a possible match for the MS/MS spectrum with protonated *m/z* 893.4624. Dataset S1, sheet four, shows the SMILES strings and delta *m/z* values for the predicted structural fragments and the observed fragments in the MS/MS spectrum. All delta *m/z* values in the table were extremely small, strongly indicating that brasilicardin A is the correct structure for this MS/MS spectrum and it matches well with the BGC identified in the genome of *Nocardia terpenica* IFM 0406 (BGC known to produce brasilicardin A, ID BGC0000632).

**Fig. S5.**
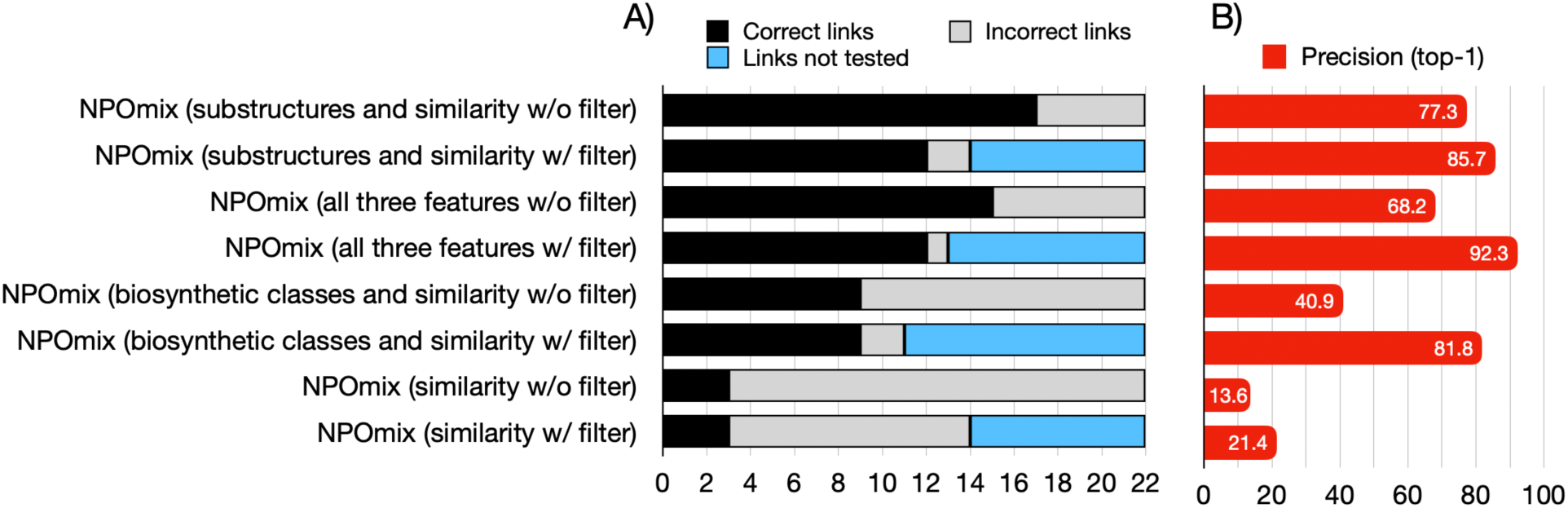
A) Histogram displaying the number of correct links (true positives, black), incorrect links (false positives, grey), links not tested (due to limitations of the tool or threshold selected, light blue) for different currently available multi-omics tools.

**Fig. S6.**
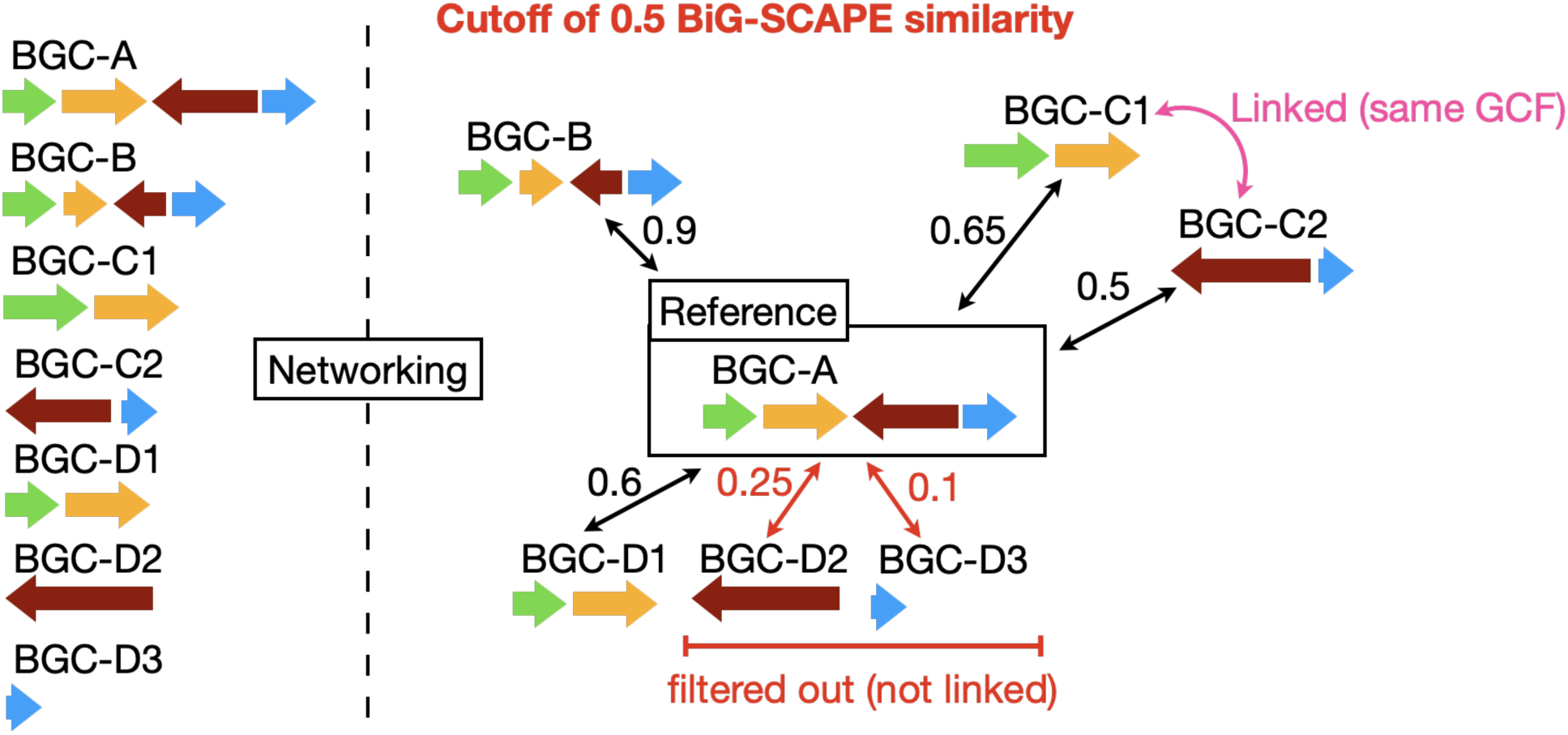
Example of fragmented biosynthetic gene clusters (BGCs) where the fragments are connected by BiG-SCAPE similarity, grouping them in a gene cluster family (in some cases). In other cases, the partial BGC still matches the reference, or some fragments are networked.

**Fig. S7.**
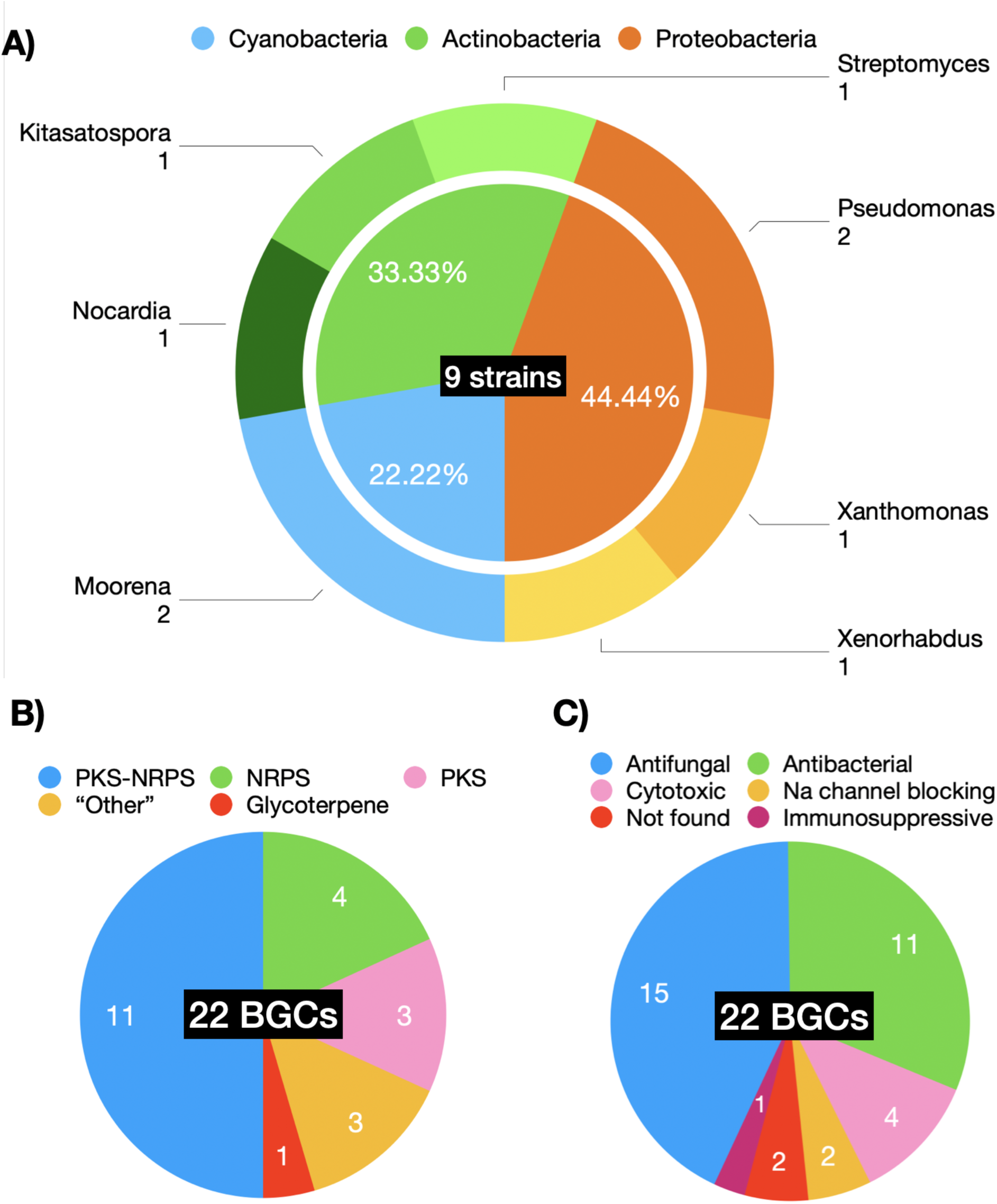
Diversity of 9 different bacteria (in A) from the PoDP database used for benchmarking NPOmix, the biosynthetic classes (in B), and the reported bioactivity (in C) found in the metabolites connected to their BGCs used in the validation dataset. Na = sodium; PKS = polyketide synthase; NRPS = nonribosomal peptide-synthetase.

**Table S1.** The number of genomes, metabolomes, total BGCs, total GCFs, MIBiG BGCs networked, GNPS metabolites dereplicated, metabolites with MIBiG BGC (and the subset correctly linked by NPOmix), and metabolites with MIBiG BGC post-filtering with 0.7 Jaccard similarity (and the subset correctly liked by NPOmix). Pal* = to palmyramide A, which the BGC is not on MIBiG yet.

| Type of feature in our dataset |  |
| --- | --- |
| Number of genomes | 1,040 |
| Number of metabolomes | 3,248 |
| Number of BGCs | 3,331 |
| Number of GCFs | 997 |
| MIBiG BGCs networked | 178 |
| GNPS metabolites dereplicated (most of them are reported at PoDP) | 50 |
| Metabolites dereplicated with MIBiG BGC | 22+Pal* |
| Metabolites dereplicated with MIBiG BGC and correctly linked by NPOmix | 17+Pal* |
| Metabolites dereplicated with MIBiG BGC post-filtering | 14 |
| Metabolites dereplicated with MIBiG BGC post-filtering and correctly linked by NPOmix | 13 |

**Table S2.**
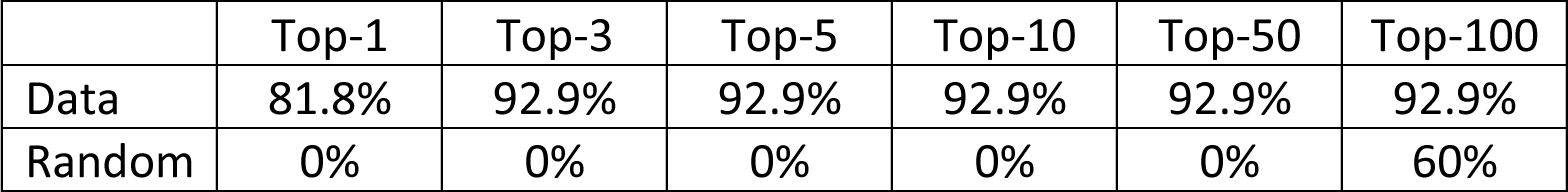
Top-*n* precision scores (how often the correct GCF label was found among the top *n* labels classified by the KNN approach) for around 14 references GNPS MS/MS spectra connected to a BGC found in the paired 1,040 (meta)genomes-MS/MS downloaded from the GNPS, MIBiG and PoDP databases (these links passed the co-occurrence threshold). These known links were obtained from the NPLinker dataset, GNPS, and PoDP databases. For this analysis, we only used similarity and biosynthetic classes as features because they are easier to annotate even from cryptic MS/MS spectra. Randomness is observed by shuffling the testing columns, experimental MS/MS names, and counting how many correct links are present between the top-*n* GCF candidates. Based on this, we believe the best performance is *n* = 3 (and it is the same value for k) for the examined dataset.

**Table S3.**
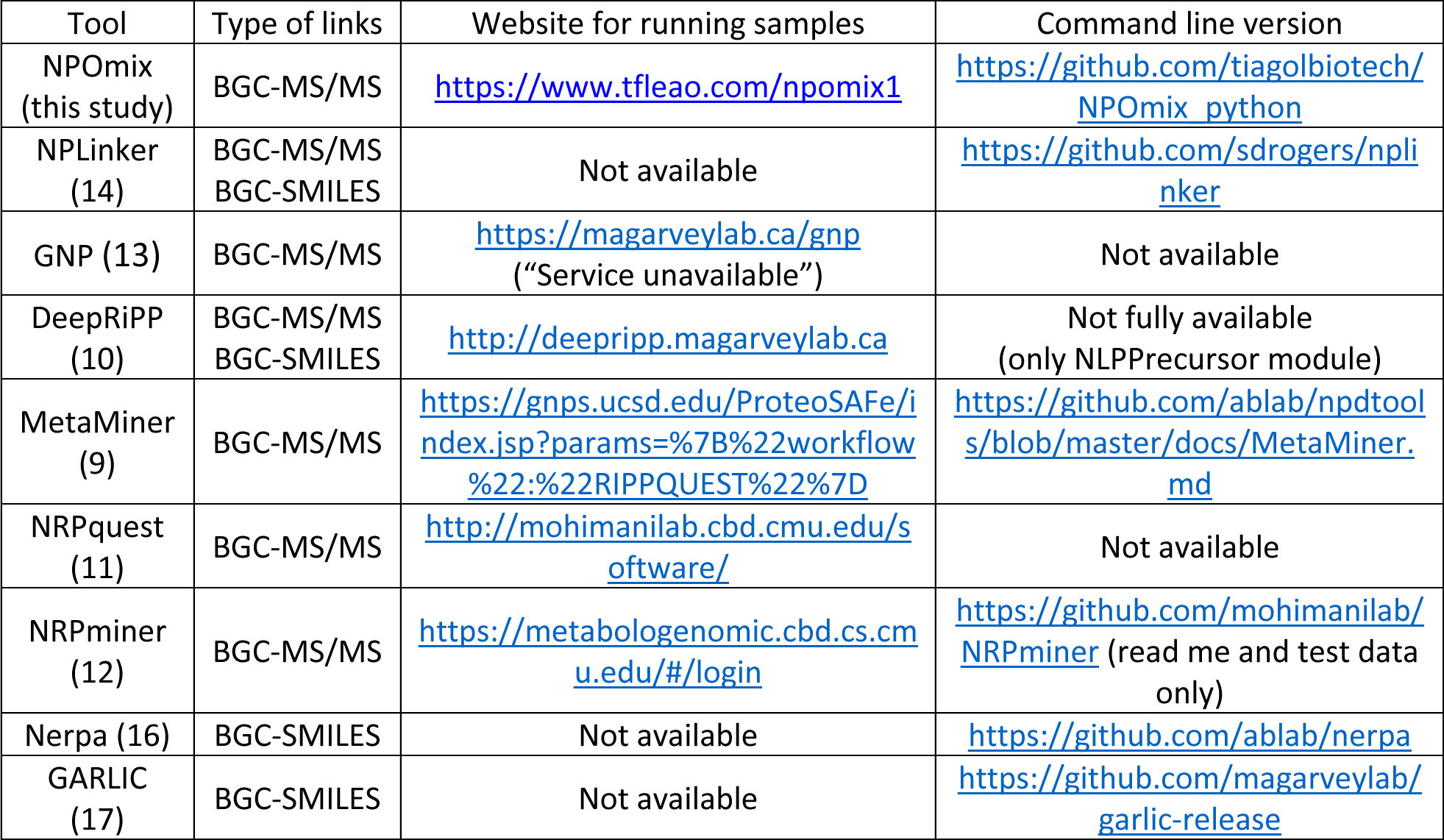
Multi-omics tools published to date, the type of links they can generate, and their websites for running samples or downloading their command-line version. Of note, only NPOmix, NPLinker, and GNP (“service unavailable”) are systematic (work for many classes of natural products). MetaMiner, and DeepRiPP are specific for ribosomally synthesized and post-translationally modified peptides (RiPPs); nonribosomal peptides (NRPs) can be processed by NRPquest (apparently discontinued), NRPminer and Nerpa. Nerpa can also process hybrid polyketides-nonribosomal peptides and GARLIC can process NRPs, polyketides, and hybrids.

